# Epigenomic Heterogeneity of Non-Functional Pancreatic Neuroendocrine Tumors Uncovered by Single nucleus and Spatial ATAC Profiling

**DOI:** 10.1101/2025.09.11.675640

**Authors:** Dejiang Wang, Xiangjun Di, Fu Gao, Gaocai Li, Liyun Lin, Shujie He, Di Zhang, Junyi Yu Jin, Yuanxin Liang, Michael Cecchini, Jill Lacy, John Kunstman, Pamela Kunz, Jing Du, Yang Liu

## Abstract

Non-functional pancreatic neuroendocrine tumors (NF-PanNETs) account for the majority of neuroendocrine neoplasms arising in the pancreas and exhibit substantial clinical and biological heterogeneity, yet their epigenetic regulation and spatial architecture remain poorly understood. Here, we present an integrative study of NF-PanNETs across multiple tumor grades using single-nucleus ATAC-seq (snATAC-seq) and spatial ATAC-seq. snATAC-seq delineates the chromatin accessibility landscapes of distinct tumor subtypes, immune cells, and cancer-associated fibroblasts (CAFs), revealing key transcription factor (TF) programs that drive tumor progression and shape microenvironmental interactions. Spatial ATAC-seq further identifies two distinct tumor-stroma ecological niches: a proliferative niche marked by MYC and FOX family, and an invasive niche enriched for Snail family TFs and KRAS pathway activity. These findings demonstrate that cellular behavior in NF-PanNETs is governed not only by intrinsic epigenetic states but also by spatial context. Together, our study provides a spatially resolved epigenomic framework for dissecting NF-PanNET heterogeneity and evolution, offering new biomarkers and regulatory axes for molecular stratification and precision therapy.

## INTRODUCTION

Pancreatic neuroendocrine tumors (PanNETs) represent the second most common type of epithelial neoplasm in the pancreas. PanNETs are defined as heterogeneous malignancies originating from the neuroendocrine precursor cells or neuroendocrine transdifferentiation of epithelial cell form pancreas^1^. They are classified into functional and non-functional (NF-PanNETs) subtypes according to the presence of clinically manifest hormonal hypersecretion syndromes^2^. NF-PanNETs account for 60%-90% of PanNETs and are increasing in incidence and prevalence over time owing to advances in molecular imaging technologies ^3^. Although some NF-PanNETs may present with a relatively indolent course, it is critical to recognize their intrinsic malignancy and variable capacity for invasion. Approximately 40%-80% of NF-PanNETs patients have distant metastasis at the time of diagnosis, most frequently involving the liver (40–93%) and exhibit significantly worse prognosis^4,5^. Currently, clinical treatment decisions are primarily guided by tumor grade and stage. Systemic therapies^6^, such as chemotherapy^7^ (e.g. temozolomide), targeted therapy^8^ (e.g. mammalian target of rapamycin (mTOR) inhibitor), hormone therapy^9,10^ (e.g. somatostatin analogs), and peptide receptor radionuclide therapy^11,12^, have showed clinical benefits on the progression-free survival, however, there is limited impact on the overall survival. Therefore, a comprehensive understanding of the molecular mechanisms driving the progression of NF-PanNETs is imperative for advancing clinical management and therapeutic strategies.

Genomic mutation landscape and dysregulated signaling pathways in NF-PanNETs have been characterized by whole-genome/exome sequencing and whole transcriptomic sequencing^13–16^. NF-PanNETs frequently harbor mutations in the DAXX/ATRX, MEN1 and mTOR pathway genes^17^. However, sporadic cases also exhibit alterations in genes such as TP53, CDKN2A, RB1, and KRAS, highlighting the molecular complexity and heterogeneity of NF-PanNETs^18,19^. While genetic mutations play a critical role in the diagnosis and treatment of NF-PanNETs, growing evidence underscores the contribution of epigenetic modifications in shaping their complex signaling landscape^20^. Epigenomic studies can provide critical insights into regulatory state transitions underlying both normal cellular functions and pathological reprogramming^21^. In particular, the advent of transposase-accessible chromatin using sequencing (ATAC-seq) has enabled the genome-wide profiling of chromatin accessibility, offering a powerful approach to map *cis*- and *trans*-regulatory activity during tumor progression^22,23^. However, bulk ATAC-seq captured an averaged chromatin accessibility profile, obscuring the genetic and regulatory heterogeneity among tumor subpopulations within the sample^23,24^. Single-cell/nucleus ATAC-seq (sc/snATAC-seq) offers novel insights into regulatory state characterization at single-cell resolution but loses spatial context during sample processing^25^. Notably, recent advances in spatial ATAC-seq now allow chromatin accessibility profiling within intact tissue architecture, restoring spatial context while retaining high-resolution epigenomic data^26,27^. This enables the mapping of epigenetic states to specific anatomical locations and microenvironmental niches, thereby improving our understanding of how spatial interactions influence cell identity, plasticity, fate decisions, and tumor-microenvironment dynamics. Therefore, extending the study from isolated single cells to multicellular hubs of interacting cells allows for more refined analyses of phenotypic heterogeneity and regulatory programs, beyond what sc/snATAC alone can reveal.

In this study, we performed an integrative chromatin accessibility analysis using snATAC-seq and spatial ATAC-seq to map the epigenetic landscape of treatment-naïve NF-PanNETs across tumor grades. Our findings reveal key transcriptional regulatory programs and spatially defined microenvironmental niches that underpin tumor progression and cellular heterogeneity, offering a framework for molecular stratification and precision therapeutic targeting.

## RESULTS

### Chromatin accessibility and cell lineage determination of NF-PanNET patient samples

We performed snATAC-seq using the Chromium platform (10x Genomics) on primary tumor specimens from nine untreated NF-PanNET patients, generating a total of 38,198 nuclei. In addition, spatial ATAC-seq was conducted on twelve NF-PanNET specimens, six of which were matched to those used for snATAC-seq (**Fig. 1a**). The snATAC-seq data were filtered using thresholds of ≥ 500 unique nuclear fragments per cell and a transcription start site (TSS) enrichment score ≥ 3 to exclude low-quality cells (**Extended Data Fig. 1a**). We identified a total of 62,896 accessible chromatin regions (ACRs) across all snATAC-seq samples, with the majority located in promoter (46.3%), intronic (29.69%), and distal intergenic (18.98%) regions.

**Fig. 1.**
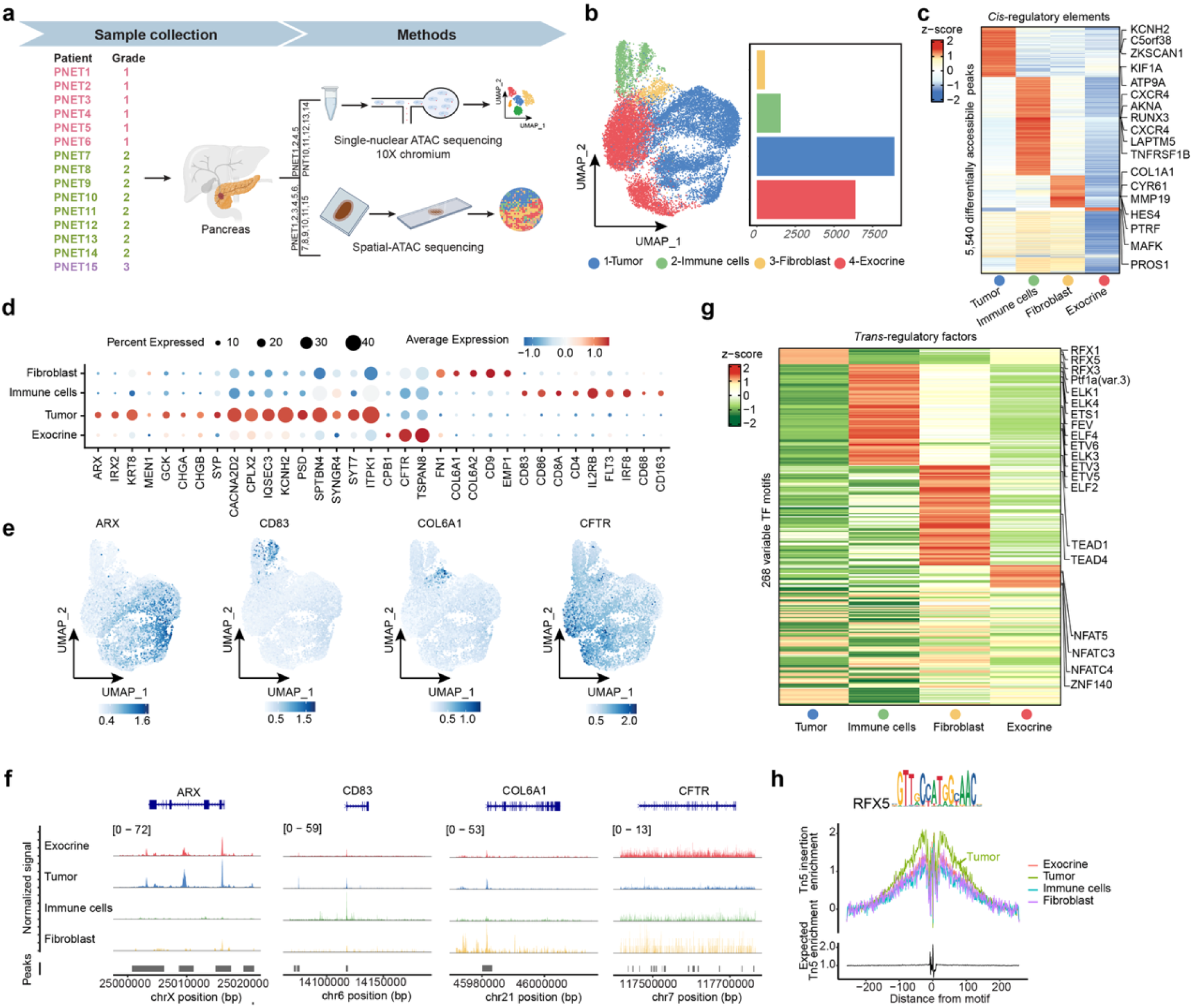
Chromatin accessibility anlaysis of cells across NF-PanNETs petients. a, Schematic overview of the study design and data generation workflow. b, UMAP of snATAC-seq profiles showing cell type annotations. Right panel: bar plot depicting the number of cells in each annotated cell subset, colored by cell type. c, Heatmap of DACRs across cell types. Color indicates z-scaled average normalized reads in each cell type. d, Dot plot of gene activity scores for representative marker genes. Dot size represents the percentage of cells with values detected in each cell type; color reflects average gene activity score. Dark red means a higher gene activity score while dark blue means a lower gene activity score. e, Spatial mapping of gene activity scores for selected marker genes across different cell types. f, Coverage plots showing chromatin accessibility at marker gene loci across cell types. g, Heatmap of differentially activated TFs per cell type. Color indicates z-scaled average TF activity score. h, Transcription factor footprinting analysis of RFX5 across cell types.

With unsupervised clusteirng, we identified four distinct cell clusters (tumor, immune, fibroblast and exocrine) based on similarities in chromatin accessibility (**Fig. 1b, Extended Data Fig. 1b**). Across these clusters, we detected 5,540 cell-type-specific differentially accessible chromatin regions (DACRs). Cell type annotation based on neighboring genes of cluster-specific *cis*-regulatory elements revealed diverse components of the tumor microenvironment (**Fig. 1c**). For instance, cluster 1 displayed accessible regions near tumor-associated genes such as KCNH2 and KIF1A. KCNH2 is frequently upregulated in pancreatic cancer compared to normal pancreatic tissue and is known to promote cell proliferation, migration, invasion, and epithelial–mesenchymal transition (EMT)^28^. KIF1A has been implicated in promoting neuroendocrine differentiation^29^. Cluster 2 was enriched for accessible chromatin regions near immune-related genes, including CXCR4, RUNX3, and LAPTM5. Cluster 3 exhibited elevated chromatin accessibility at loci neighboring fibroblast marker genes such as COL1A1 and MMP19, suggesting a fibroblast identity. Next, the gene activity score matrix was generated using Cicero^30^, and cell type annotation was performed based on the gene activity scores of canonical marker genes (**Fig. 1d**). For instance, tumor cells were marked by CHGA, CHGB, CACNA2D2, and SYT7, whereas fibroblasts were identified by FN1, COL6A1, and COL6A2. Immune cells were annotated based on high gene activity scores of CD86, CD4, and IL2RB, and exocrine cells were defined by elevated scores of CFTR and TSPAN8^31^. We further projected cell type specific gene activity scores onto the snATAC-seq profiles, which supported the identities of clusters defined by *cis*-element accessibility (**Fig. 1e, f**).

Next, we assessed chromatin accessibility at *cis*-regulatory elements harboring TF binding motifs using chromVAR^32^. We observed increased TF deviation scores for the RFX family, known to contribute to tumor aggressiveness^33^ in the tumor cluster (**Fig. 1g, Extended Data Fig. 1e**). In line with this, TF footprinting analysis revealed enhanced chromatin accessibility surrounding RFX binding sites in tumor cells (**Fig. 1h**), consistent with elevated RFX activity. Likewise, deviation scores for the ETS family, including crucial regulators of T cells, B cells, macrophages, neutrophils, and dendritic cells^34^ were specifically elevated in immune cell clusters (**Fig. 1g**). Analysis of peak genomic distribution and 15-core chromHMM states indicated that peaks in immune cells and fibroblasts were predominantly located at promoters and enhancers, highlighting the active regulatory landscape within the tumor microenvironment (**Extended Data Fig. 1c, 1d**).

### Characterization of chromatin accessibility alterations across tumor grades

To investigate chromatin accessibility dynamics across tumor grades in NF-PanNETs, we isolated tumor cells from snATAC-seq data. Using latent semantic indexing (LSI) and UMAP for dimensionality reduction and visualization, we identified nine tumor subclusters (Tumor_1 to Tumor_9; **Fig. 2a & Extended Data Fig. 2a**), each defined by distinct chromatin accessibility profiles and gene activity signatures (**Fig. 2b**, **2c**). Notably, the distribution of these subtypes differed markedly between tumor grades. Grade 1 tumors predominantly comprised Tumor_1 to Tumor_4, while Grade 2 tumors included Tumor_5 to Tumor_9. Previous classifications of neuroendocrine carcinomas have identified five molecular subtypes defined by the expression of ASCL1, NEUROD1, HNF4A, POU2F3, and YAP1^35^. In our cohort, Grade 1 tumor cells displayed elevated gene activity scores for HNF4A, while POU2F3 was specifically enriched in Tumor_5, a Grade 2 subtype. Other Grade 2 subtypes lacked canonical subtype markers but instead showed increased activity of genes such as PCSK1, PTEN, TRIM24, and TRIM28, indicating additional regulatory diversity beyond classical subtype definitions.

**Fig. 2.**
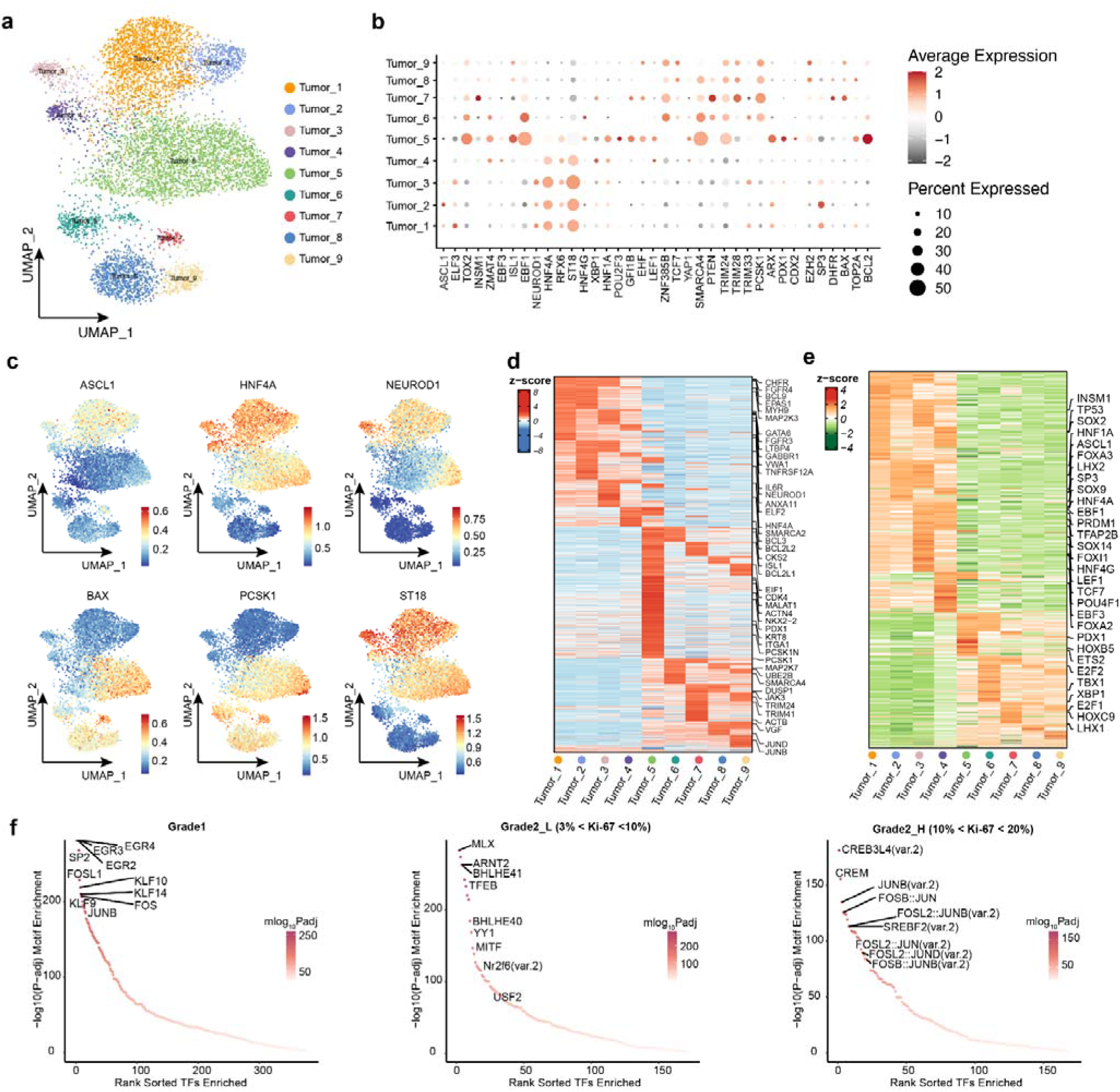
Tumor heterogeneity characterized by chromatin accessibility dynamics across tumor grades. a, UMAP projection of annotated tumor cells based on snATAC-seq profiles. b, Dot plot illustrating gene activity scores of representative marker genes across tumor subtypes. Dot size indicates the percentage of cells with detectable gene activity; color reflects the average activity score. Dark red means higher gene activity score, dark gray means lower gene activity score. c, Spatial mapping of gene activity scores for selected subtype-specific marker genes. d, Top DACRs identified for each tumor subtype. Color indicates the z-scaled average normalized read counts per tumor. e, Differentially activated TFs across tumor subtypes, with color representing z-scaled average TF activity scores. f, Enriched TF motifs associated with tumor grade-specific DACRs.

Next, we performed pathway enrichment analysis, which revealed that tumor subpopulations were enriched in biological processes including cell proliferation, epithelial-to-mesenchymal transition (EMT), and immune-related pathways (**Extended Data Fig. 2b**). Tumor_4, a Grade 1 cluster, was enriched for proliferation-related gene sets, including E2F targets, G2M checkpoint, and MYC targets. Conversely, Tumor_1 and Tumor_5, which exhibited lower proliferative activity, showed enrichment in pathways such as KRAS signaling up and inflammatory response, both implicated in pancreatitis and pancreatic intraepithelial neoplasia (PanIN)^36^. DACRs across subtypes and grades further underscored the epigenetic heterogeneity of NF-PanNETs (**Fig. 2d & Extended Data Fig. 2c**). Genomic annotation and ChromHMM state analysis of DACRs revealed distinct distributions between tumor grades, particularly in promoter and enhancer regions (**Extended Data Fig. 2d, 2e**).

We next examined TF activity across tumor subtypes using chromVAR-derived deviation scores (**Fig. 2e**). HNF1A, HNF4A, and HNF4G showed higher activity in Grade 1 tumors, indicating well-differentiated neuroendocrine states. Whereas Grade 2 tumors were characterized by increased activity of ETS2, E2F2, and E2F1, reflecting dedifferentiated, proliferative, and aggressive states. Footprinting analysis confirmed greater chromatin accessibility at HNF1A binding sites in Grade 1 tumors compared to Grade 2 (**Extended Data Fig. 2f**). To determine whether these accessibility changes reflect altered transcriptional regulation, we performed motif enrichment analysis on DACRs (**Fig. 2f & Extended Data Fig. 2g**). Motifs for KLF and EGR family members were enriched in Grade 1, while TFEB and YY1 motifs predominated in low-proliferative Grade 2 tumors (Grade2_L, 3%< Ki-67 <10%). In contrast, AP-1 motifs were significantly enriched in highly proliferative Grade 2 tumors (Grade2_H, 10%< Ki-67 <20%). Given its known role in regulating genes involved in proliferation, differentiation, apoptosis, angiogenesis, and invasion^37^, the enrichment of AP-1 highlights the aggressive regulatory landscape in Grade2_H tumors.

### Dynamic landscape of transcription factors associated with tumor progression

Having identified tumor-subtype-specific TFs associated with different histological grades, we hypothesized that these TFs might constitute a coordinated regulatory program governing tumor initiation and progression. To investigate the dynamic changes in regulatory states during NF-PanNET evolution, we performed trajectory inference using all tumor cells. As shown in **Fig. 3a**, the reconstructed pseudotime trajectory revealed a continuous evolutionary path, with Grade 1 tumor cells positioned at the origin and Grade 2 tumor cells located toward the terminal end, consistent with their pathological progression.

**Fig. 3.**
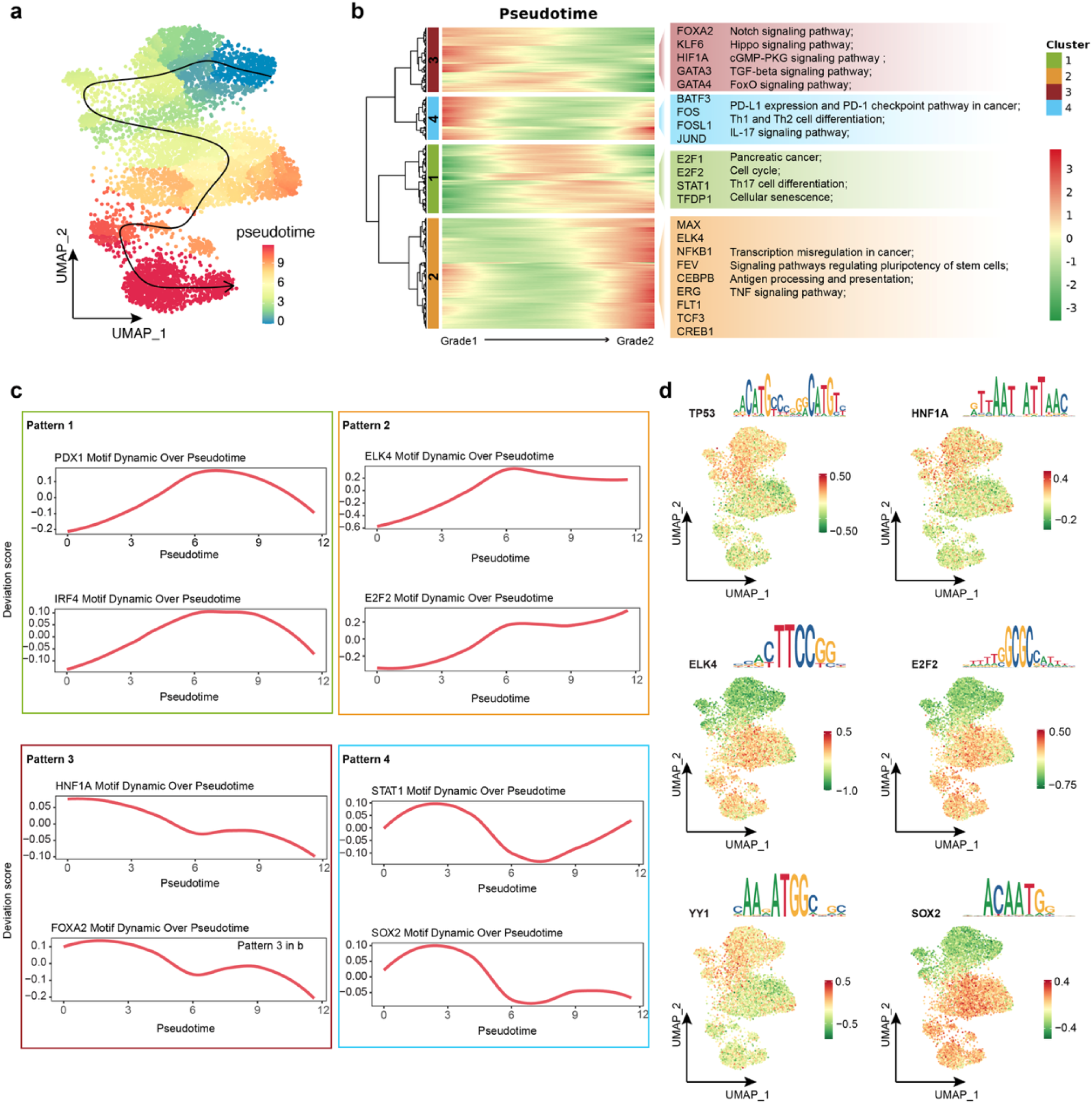
Dynamic chromatin accessibility and transcription factor activity changes during tumor progression. a, Pseudotime trajectory of tumor cell differentiation inferred from snATAC-seq profiles. b, Heatmap showing dynamic changes in TF activity along differentiation trajectories. Right panel highlights representative TFs and the enriched functional pathways. c, Line plots displaying the dynamic trends of TF activity along pseudotime during tumor progression. d, UMAP distribution of TF deviation scores for selected key TFs across different tumor grades.

We next examined the dynamic changes in TF activity along the pseudotime trajectory by calculating TF deviation scores across the continuum of tumor cell evolution. Unsupervised clustering of the smoothed scores resolved four temporal patterns (**Fig. 3b**, **3c**). Patterns 1–2 comprised TFs whose activity increased monotonically along the trajectory (e.g., ELK4, E2F2), with target enrichment in cell-cycle, MAPK/ETS, Th17 differentiation, TNF signaling, and transcriptional misregulation in cancer, consistent with progressive proliferative/inflammatory rewiring. Conversely, Pattern 3 contained TFs decreasing with pseudotime (e.g., FOXA2, HNF1A), whose targets were enriched for Notch, Hippo, cGMP–PKG, TGF-β, and FoxO pathways, programs prominent in early tumor states. Interestingly, TFs in Pattern 4 displayed a distinct trajectory: their activity remained relatively stable during early and late stages but showed significant downregulation during the intermediate stage (**Fig. 3c**). Functional annotation of these TFs revealed enrichment in immune-related pathways, including PD-L1 expression and PD-1 checkpoint, Th1/Th2 differentiation, and IL-17 signaling, underscoring the transient reprogramming of the immune microenvironment and suggesting potential windows for immunotherapeutic intervention. We highlighted representative TFs critical for tumor progression and mapped their activity across pseudotime (**Fig. 3c**). For instance, HNF1A, and FOXA2 exhibited high activity early in the trajectory, while ETS family members became active later, consistent with the emergence of an immune-modulating microenvironment. To validate these findings, we projected TF deviation scores onto the UMAP of snATAC-seq profiles (**Fig. 3d**). Immune checkpoint regulators such as TP53 were predominantly active in Grade 1 tumors, whereas TFs including ELK4, E2F2, and SOX2 were enriched in Grade 2, supporting their roles in tumor progression and grade-specific regulatory states.

### Chromatin landscape of intratumoral immunity

To characterize the immunosuppressive environment in NF-PanNETs, we subdivided immune cells into myeloid and lymphoid compartments (**Fig. 4a**, **4b, Extended Data Fig. 3a**). We classified myeloid cells by their high gene activity scores for ITGAM, CD68, and CD163. Within the lymphocyte compartment, distinct subsets were identified, including T cells (CD8A, CD8B), B cells (CD19, CD79A), and natural killer (NK) cells (SPON2, IL2RB, KLRD1). NK cells, as key players in cancer immunosurveillance, exert anti-tumor functions through natural cytotoxicity and antibody-dependent cellular cytotoxicity^38^. KEGG pathway enrichment analysis revealed that NK cells in NF-PanNETs were functionally enriched for pathways including NK cell-mediated cytotoxicity, Fc gamma R–mediated phagocytosis, antigen processing and presentation, and apoptosis (**Fig. 4c**). Furthermore, NK cells displayed increased chromatin accessibility at the promoter region of the IL2RB locus (**Extended Data Fig. 3b**). In contrast, B cells contribute to adaptive immunity primarily through antigen presentation and antibody secretion^39^. KEGG functional annotation analysis indicated enrichment of gene sets associated with the B cell receptor signaling pathway, NF-κB signaling, and cellular senescence.

**Fig. 4.**
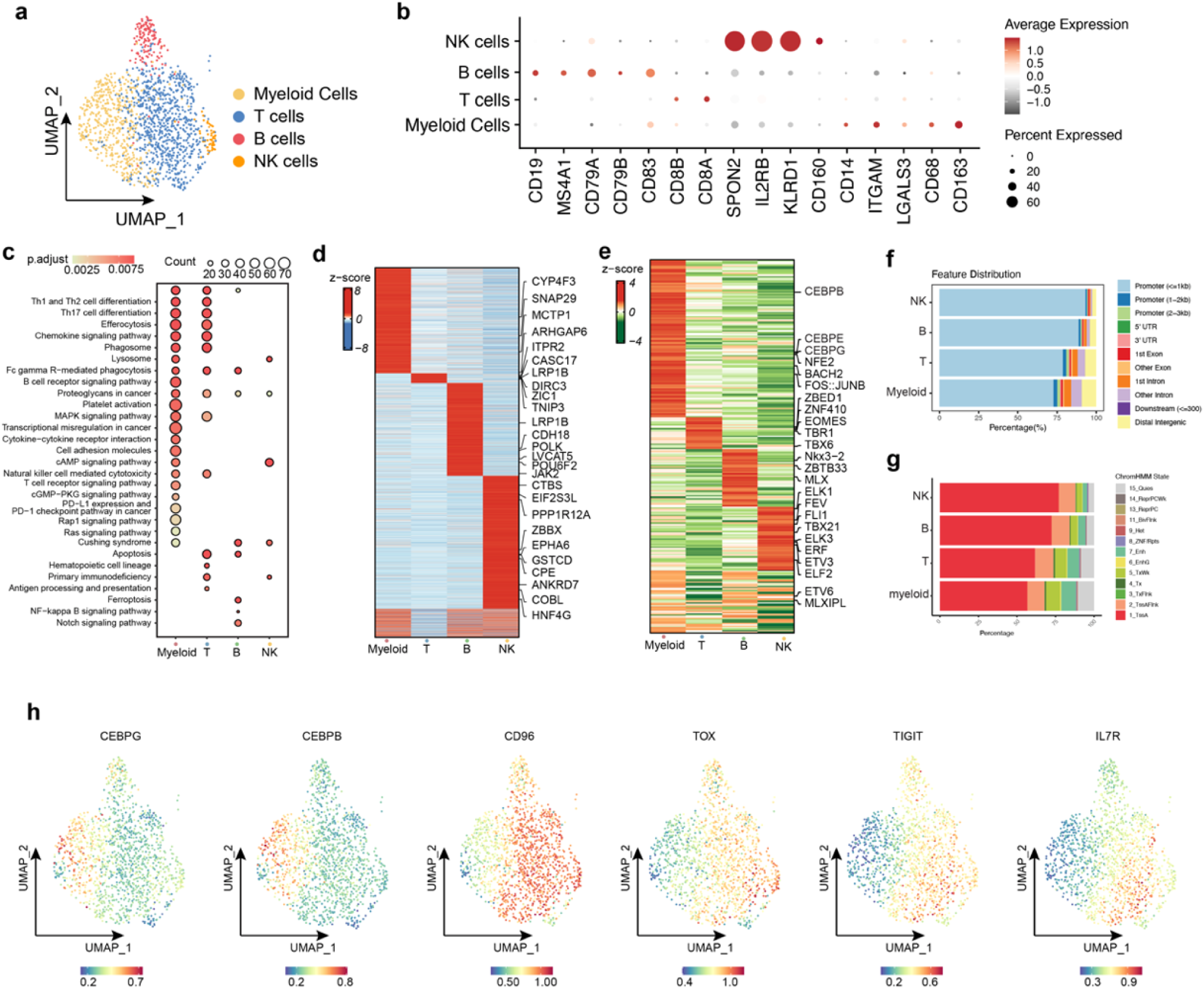
Chromatin landscape of intratumoral immunity in NF-PanNETs. a, UMAP projection of snATAC-seq profiles showing annotated immune cell subtypes. b, Dot plot of gene activity scores for canonical marker genes across immune cell subtypes. Dot size represents the proportion of cells expressing the gene; color indicates the average gene activity score. Dark red means higher gene activity score, dark gray means lower gene activity score. c, Immunologic signature and KEGG pathway enrichment analysis of DACRs in each immune cell subtype. d, Heatmap showing the top DACRs for each immune cell subtype. Color scale reflects z-scored average normalized accessibility per subtype. e, Heatmap of TF activity scores across immune cell subtypes. Color indicates z-scored deviation scores. f, Genomic feature distribution of subtype-specific DACRs, including promoter, enhancer, and intergenic regions. g, ChromHMM state annotation of DAPs for each immune cell subtype. State definitions: TssA (Active TSS), TssAFlnk (Flanking Active TSS), TxFlnk (Transcription at gene 5′/3′), Tx (Strong transcription), TxWk (Weak transcription), EnhG (Genic enhancers), Enh (Enhancers), ZNF/Rpts (ZNF genes & repeats), Het (Heterochromatin), TssBiv (Bivalent/poised TSS), BivFlnk (Flanking bivalent TSS/Enhancer), EnhBiv (Bivalent enhancer), ReprPC (Repressed PolyComb), ReprPCWk (Weak repressed PolyComb), Quies (Quiescent/Low). h, UMAP mapping of gene activity scores at representative marker loci across immune subtypes.

Compared with NK and B cells, myeloid cells exhibited broader involvement in multiple signaling pathways critical to immune regulation and tumor progression^40^. Pathway analysis revealed that myeloid cells were enriched in chemokine signaling, MAPK signaling, cytokine–cytokine receptor interaction, cAMP signaling, cGMP–PKG signaling, Rap1 signaling, and Ras signaling pathways (**Fig. 4c**), underscoring their central role in shaping the TME. Consistently, differential chromatin accessibility at *cis*-regulatory elements revealed increased accessibility in key genomic regions within myeloid cells (**Fig. 4d**). Notable examples included CYP4F3, an enzyme involved in leukotriene B4 degradation and inflammation resolution^41^, and SNAP29, a gene implicated in vesicle fusion and immune regulation^42^. Myeloid-derived suppressor cells (MDSCs), which arise in contexts of chronic inflammation and advanced cancers, are known to drive immune evasion and tumor progression^43^. MDSC-like cells are generally characterized by a two-signal process: the first involves STAT3 activation and regulation by transcription factors such as CEBPA, CEBPB, and IRF8^44,45^; the second engages the NF-κB pathway, stimulated by inflammatory mediators like TNF-α, various interleukins, and HMGB1^46,47^. In our data, we observed increased chromatin accessibility and gene activity scores for CEBPB, CEBPE, and CEBPG in myeloid cells, supporting the presence of MDSC-like populations in NF-PanNETs (**Fig. 4e, 4h**). Genomic annotation and chromHMM segmentation of accessible chromatin region further revealed that myeloid and T cell peaks were less enriched at promoter and enhancer regions compared to other immune subtypes (**Fig. 4f, 4g**), reflecting a transcriptionally suppressed or altered regulatory state characteristic of immune dysfunction in the TME.

To further dissect myeloid heterogeneity, we reclustered myeloid cells into four subsets (**Fig. 5a**). We then assessed the expression of markers associated with M1, M2a, M2c, and tumor-associated macrophages (TAMs) (**Extended Data Fig. 3c**). Interestingly, none of the four macrophage subsets conformed strictly to canonical M1/M2 polarization, suggesting more complex or intermediate activation states. Projection of gene activity scores for MDSC-associated genes revealed that macrophage subset 1 was enriched for IRF8 and ZBTB46, both associated with early MDSC-like differentiation (“first-signal”), while macrophage subset 3 expressed HMGB1 and CCL18, indicative of inflammatory activation and terminal suppression (“second-signal”) (**Fig. 5b**). Pathway enrichment analysis confirmed distinct functional profiles among the subsets, with macrophages 1 and 3 showing strong enrichment for hallmark gene sets including TNF-α signaling via NF-κB, apoptosis, the p53 pathway, and IL2–STAT5 signaling (**Fig. 5c**). To uncover the transcriptional regulatory programs underpinning each macrophage subtype, we calculated TF deviation scores and identified subtype-specific TFs (**Fig. 5d**). Macrophage 0 was characterized by enrichment of SREBF1 and SREBF2, whereas macrophage 1 exhibited elevated activity of POU1F1, POU3F1, and POU3F2. In contrast, macrophage 3 showed specific enrichment of the HOX and TEAD transcription factor families. These findings highlight the functional diversity of macrophage subsets in the immunosuppressive TME of NF-PanNETs and suggest distinct roles in modulating immune escape and tumor progression.

**Fig. 5.**
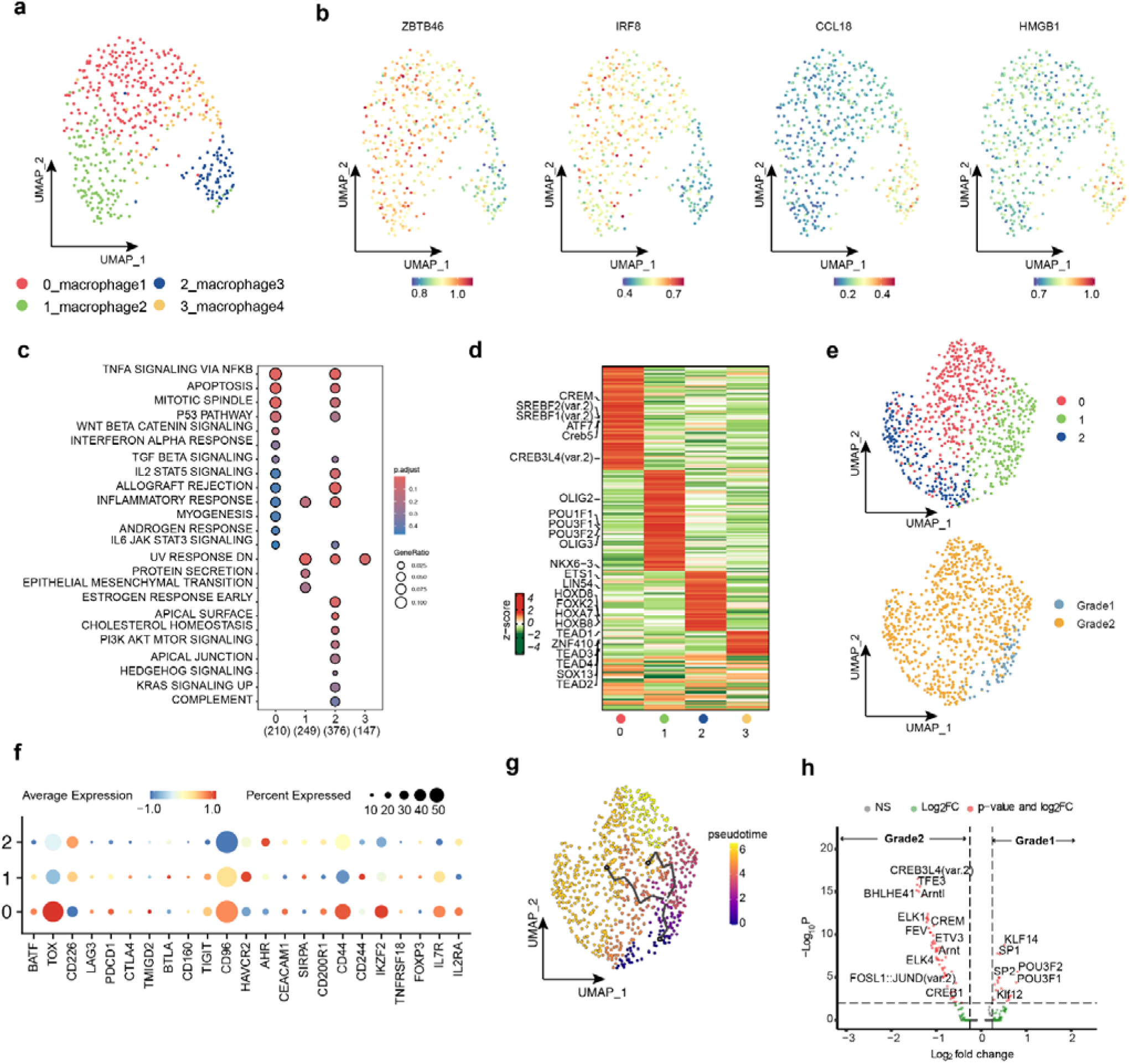
Epigenetic heterogeneity of macrophages and T cells in NF-PanNETs. a, UMAP projection showing annotated snATAC-seq profiles of macrophage subtypes. b, UMAP mapping of gene activity scores for representative marker genes across macrophage subtypes. c, Enriched hallmark pathways based on DACRs upregulated in each macrophage subtype. Dot size reflects gene ratio; color indicates adjusted adjusted *p*-value. d, Differentially activated TFs across macrophage subtypes. Colors represent z-scaled average TF activity scores e, UMAP projection showing annotated snATAC-seq profiles of T cell subtypes. f, Dot plot of gene activity scores for marker genes of T cell subtypes. Dot size indicates the percentage of cells expressing the gene; color represents average activity score. Dark red means higher gene activity score, dark blue means lower gene activity score. g, Pseudotime trajectory of T cell differentiation. h, Volcano plot showing differentially accessible TF motifs between Grade 1 and Grade 2 NF-PanNETs.

To delineate the heterogeneity of T cells in the TME of NF-PanNETs, we selected cells expressing canonical T cell markers CD8A and CD8B, identifying three transcriptionally distinct clusters (**Fig. 4b & Fig. 5e**). Clusters 0 and 2 were predominantly enriched in Grade 2 tumors, whereas Cluster 1 was the major subset found in Grade 1 tumors (**Fig. 5e**), suggesting a potential grade-specific T cell distribution.

To further characterize these subsets, we examined gene activity scores of T cell subtype markers and immune regulatory genes (**Fig. 5f**). Notably, Cluster 0 displayed elevated gene activity scores for TOX, CD96, CD44, and IKZF2, indicating a phenotype consistent with “exhausted-like” T cells. Functional enrichment analysis of Cluster 0 further supported this interpretation, with significant enrichment of the Hedgehog signaling pathway (**Extended Data Fig. 3d & 3e**), a pathway known to promote immune suppression and foster an immunosuppressive microenvironment^48^. However, the exhausted phenotype of Cluster 0 did not align with classical exhaustion signatures. Specifically, key inhibitory receptors such *as* PDCD1 (PD-1) and CTLA4 were not prominently expressed. This observation is consistent with prior findings that PD-1/PD-L1 expression is generally low or absent in small intestinal and pancreatic neuroendocrine tumors^49^. These findings suggest the presence of alternative, non-canonical immunosuppressive mechanisms in the NF-PanNET microenvironment. Moreover, we observed higher gene activity scores for several non-classical immune checkpoint suppressors in Grade 2 tumors (**Extended Data Fig. 3e**).

To further explore potential functional transitions, we performed pseudotime trajectory analysis using Monocle3^50–53^, which revealed a progression from T cells in Grade 1 tumors toward those in Grade 2 (**Fig. 5g**). This trajectory suggests a dynamic change in T cell state associated with tumor progression. We next investigated the transcriptional regulatory landscape underlying these differences by comparing motif enrichment between Grade 1 and Grade 2 T cells (**Fig. 5h**). In Grade 2 tumors, we observed significant enrichment of TFE3 and CREM motifs. TFE3, in coordination with TFEB, modulates gene expression in T cells, particularly under stress conditions^54^. CREM plays a critical role in regulating T cell tolerance and exhaustion^55^, supporting the presence of “exhausted-like” T cells in Grade 2. In contrast, T cells in Grade 1 tumors exhibited increased accessibility of POU family motifs, which are linked to cell identity and stem-like transcriptional programs^56^. Collectively, these results suggest that T cells in Grade 2 tumors adopt a more stress-adapted and exhausted regulatory state, whereas those in Grade 1 maintain a quiescent or stem-like chromatin configuration. This shift in transcriptional control may contribute to the immunosuppressive and progressive nature of higher-grade NF-PanNETs.

### Chromatin landscape of the stromal microenvironment

CAFs are pivotal components of the tumor microenvironment and key drivers of tumor progression in various solid malignancies^57,58^. Unsupervised clustering identified five distinct CAF subpopulations (clusters 0–4), each exhibiting unique chromatin accessibility landscapes (**Fig. 6a, b**). Gene activity scores of canonical CAF markers were computed to annotate these clusters (**Fig. 6c, Extended Data Fig. 4a**). Clusters 0 and 1 were identified as classical myofibroblastic CAFs (myCAFs), characterized by elevated activity scores of ACTA2, TAGLN, FN1, and FAP—markers indicative of contractility and ECM deposition, which contribute to tumor fibrosis and increased tissue stiffness^57^. Notably, pathway enrichment analysis indicated enrichment of EMT programs in CAFs from clusters 1 and 2 (**Fig. 6g**). Cluster 3 displayed an immunomodulatory phenotype, with high activity of CD74 and THY1, suggestive of antigen-presenting CAFs (apCAFs) potentially involved in modulating immune responses^59^. In contrast, cluster 2 exhibited a mechanobiological signature, marked by co-activation of DPT and TPM1, implying a role in balancing matrix organization and cellular contractility. Cluster 4 represented a matrix-remodeling subtype, with elevated gene activity of MMP11 and the transcriptional regulator HOPX, suggesting active ECM degradation and promotion of a pro-invasive niche^60^. To understand the regulatory architecture underlying CAF diversity, we assessed both differential chromatin accessibility and TF activity across subtypes (**Fig. 6d, e**). Each cluster exhibited a unique set of DACRs, reflecting subtype-specific regulatory programs. For instance, cluster 0 showed increased accessibility near metabolic genes such as PDK4, indicating a metabolic reprogramming signature. Cluster 3 exhibited open chromatin near DLL4, a Notch signaling ligand associated with angiogenic regulation^61^. Cluster 4 showed accessible regions near genes related to cell motility and cytoskeletal remodeling, including CD164 and ARHGAP19, reinforcing its invasive phenotype.

**Fig. 6.**
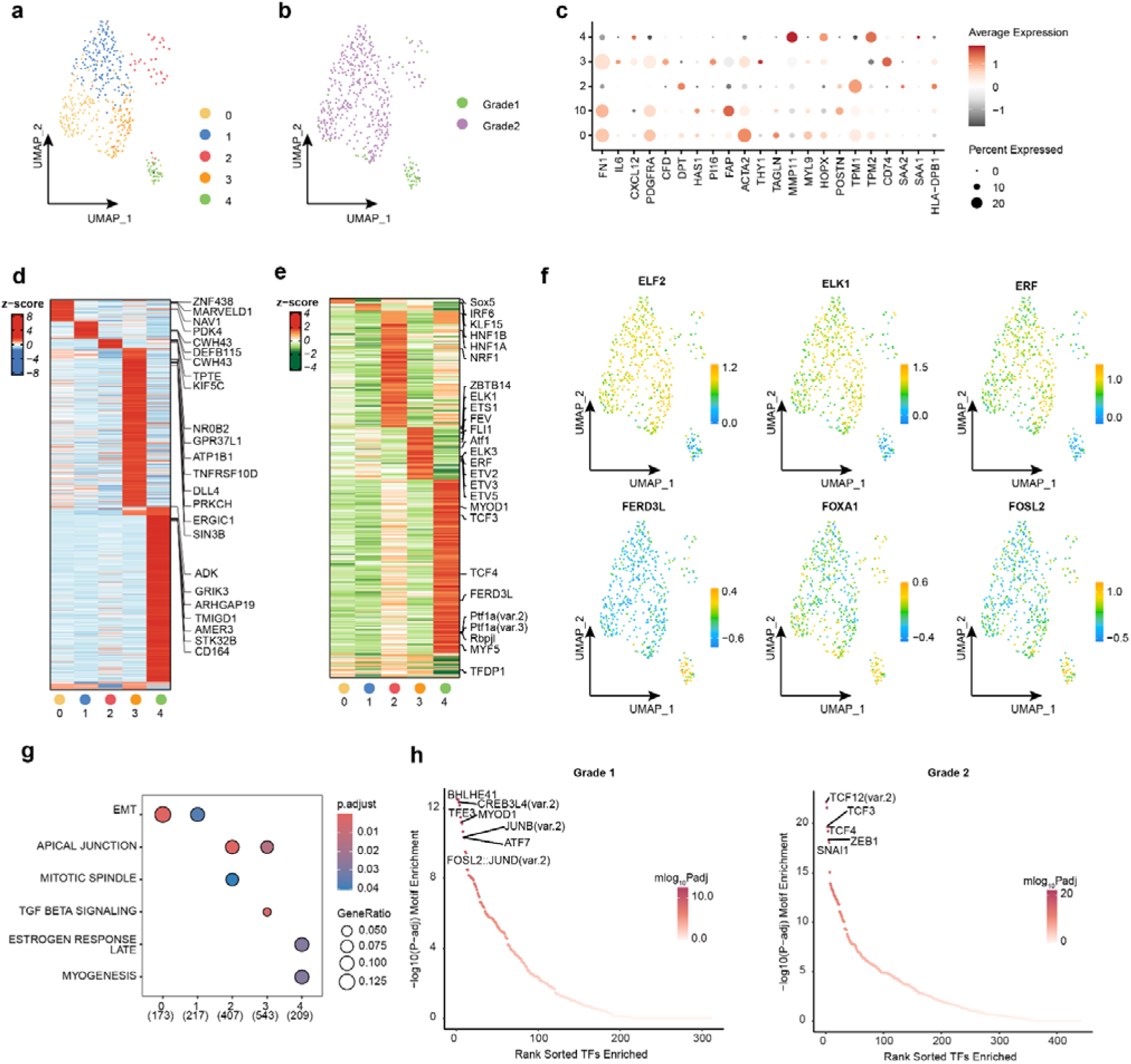
Chromatin landscape of the stroma microenvironment. a, UMAP projection of snATAC-seq profiles showing annotated subtypes of CAFs. b, UMAP projection of CAFs colored by tumor grade. c, Dot plot showing gene activity scores of representative marker genes across CAF subtypes. Dot size represents the percentage of cells expressing the gene; color indicates the average gene activity score. Dark red means higher gene activity score, dark gray means lower gene activity score. d, Heatmap of the top DACRs for each CAF subtype. Color represents z-scaled average normalized accessibility. e, Heatmap of differentially activated TFs across CAF subtypes. Color represents z-scaled average TF activity score. f, UMAP mapping of selected TF deviation scores in CAFs across the tissue. g, Significant hallmark pathway enrichments of DACRs upregulated in each CAF subtype. Bubble size represents gene ratio, and color indicates adjusted p value. h, Enrichment of CAF-related motifs within tumor grade-specific DACRs.

To further investigate the regulatory changes across tumor grades, we performed motif enrichment analysis on grade-specific DACRs within CAFs. This revealed striking reprogramming of transcriptional regulatory networks in high-grade tumors (**Fig. 6h**). In Grade 2, we observed strong enrichment of motifs for pro-metastatic TFs such as SNAI1, ZEB1, and members of the TCF family (e.g., TCF4, TCF12)—key orchestrators of EMT and invasion. These findings provide a direct link between chromatin accessibility and the aggressive behavior of CAFs in high-grade tumors. In contrast, Grade 1 CAFs were enriched for motifs associated with stress response, differentiation, and moderate proliferative programs, including FOSL2/JUNB (AP-1 complex), CREB3L1, and BHLHE41 (**Fig. 6h**). These TFs reflect a less aggressive transcriptional landscape, consistent with the lower invasive potential of early-stage disease. Collectively, these results highlight profound epigenetic heterogeneity within CAFs and underscore a regulatory shift toward pro-invasive, EMT-associated programs as tumor grade increases. This chromatin remodeling likely contributes to the enhanced malignancy and immunosuppressive microenvironment characteristic of high-grade NF-PanNETs.

### Spatial chromatin accessibility dynamics underpin NF-PanNET tumor progression

Spatial ATAC-seq enables the mapping of chromatin accessibility within the native tissue architecture, complementing insights from snATAC-seq. We performed spatial ATAC-seq on tissues of 12 primary NF-PanNETs (**Extended Data Fig. 5-6**). H&E stained tissue sections were annotated by pathologists (**Extended Data Fig. 7**). Integration of spatial ATAC-seq data from 12 NF-PanNET samples revealed six distinct clusters defined by chromatin accessibility profiles, along with 93,119 DACRs (**Fig. 7a, Extended Data Fig. 9a**). Tumors classified as Grade 1 and Grade 2 primarily comprised five clusters (c0-c4), whereas c5 appeared to represent a malignant population unique to Grade 3 tumors (**Fig. 7a-7b, red circle**). Cluster-specific cis-elements and their nearby genes were used to annotate the clusters, revealing a distinct chromatin accessibility signature in Grade 3 tumors (**Fig. 7c**). Mapping c5 back onto the tissue sections showed that its spatial distribution corresponded to the tumor region of sample PNET5, consistent with H&E staining (**Fig. 7d, Extended Data Fig. 7**I).

**Fig. 7.**
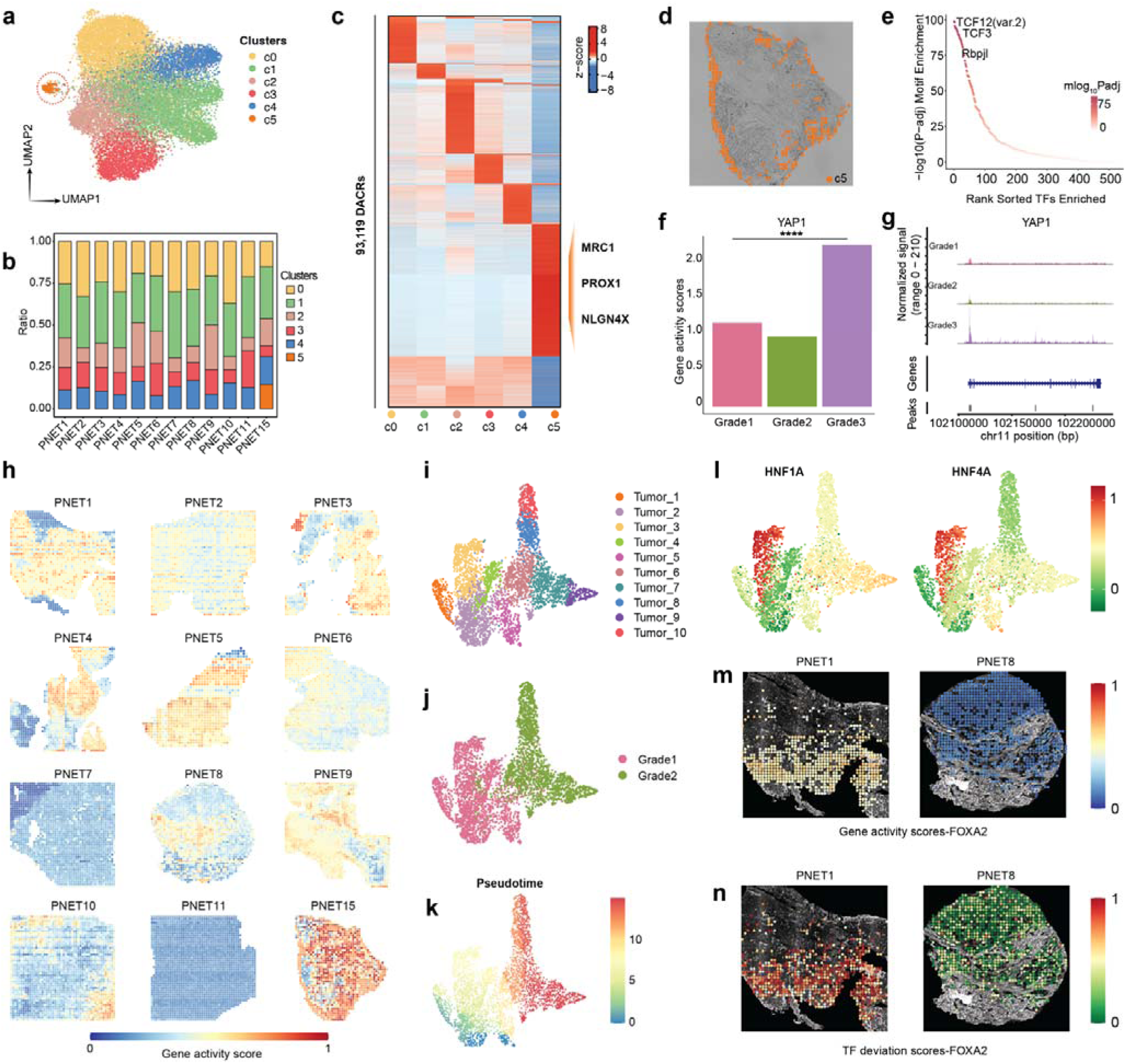
Spatial chromatin accessibility landscape of cells across twelve NF-PanNETs. a, UMAP projection of spatial-ATAC-seq profiles showing annotated cell types. A red circle highlights the unique cluster c5 identified in a Grade 3 tumor. b, Bar plot displaying the proportion of each cell type in individual samples. c, Heatmap of top DACRs specific to Grade 3 tumors. Colors represent z-scaled average normalized read counts across cell types. d, Spatial localization of cluster c5 within sample PNET5 **e,** Enriched TF motifs identified in sample PNET15. **f,** Bar plot showing YAP1 gene activity scores across tumor grades. **g,** Genome browser coverage plot illustrating chromatin accessibility at the YAP1 locus across tumor grades. **h,** Spatial distribution of YAP1 gene activity scores across representative samples. **i–j,** UMAP projections of spatial-ATAC-seq profiles for tumor cells: **i,** grouped by cell identity; **j,** stratified by tumor grade. **k,** Pseudotime trajectory representing tumor cell differentiation states. **l,** UMAP projections of TF deviation scores for HNF1A and HNF4A. **m,** Spatial maps of FOXA2 gene activity scores in samples PNET1 and PNET8. **n,** Spatial distribution of FOXA2 TF deviation scores in PNET1 and PNET8.

Subsequently, we focused on Grade 3 tumors and identified three major cell-type clusters, tumor cells, immune cells, and fibroblast cells (**Extended Data Fig. 8j**). Motif enrichment analysis revealed overrepresentation of TCF3 and TCF12 motifs in sample PNET15 (**Fig. 7e**). These transcription factors are known regulators of oncogenic processes including EMT, tumor invasion, and stemness, and have been implicated in solid tumors such as pancreatic cancer, glioblastoma, and breast cancer, further confirming the link between chromatin accessibility and the aggressive behavior in high-grade tumors^62,63^. We further calculated gene activity scores and found that YAP1 exhibited significantly higher activity in Grade 3 tumors compared to Grade 1 and 2 (**Fig. 7f, 7h**). YAP1, a downstream effector of the Hippo signaling pathway, is involved in tissue repair, regeneration, and tumorigenesis, and is recognized as a prognostic biomarker in pancreatic cancer. It has also been linked to the remodeling of the extracellular matrix^64^. Browser track visualization confirmed increased chromatin accessibility at the YAP1 locus in Grade 3 tumors (**Fig. 7g**). Taken together, these findings demonstrate that Grade 3 tumors possess a distinct and more aggressive chromatin accessibility landscape, with molecular features similar to pancreatic cancer^35^, distinguishing them from Grade 1 and Grade 2 NF-PanNETs.

Next, we integrated tumor clusters from Grade 1 and Grade 2 NF-PanNETs and identified 10 subclusters, designated Tumor-1 to Tumor-10 (**Fig. 7i, 7j**). Among these, Grade 1 tumors were composed of Tumor_1 to Tumor_5, while Grade 2 tumors consisted of Tumor_6 to Tumor_10. TF deviation scores revealed that HNF1A, HNF4A, and HNF4G were significantly enriched in Grade 1 tumors, consistent with the results from snATAC-seq analysis (**Fig. 7l, Extended Data Fig. 9b-9c**). To further explore the evolutionary trajectory during NF-PanNET progression, we performed pseudotime trajectory analysis using spatial ATAC-seq data. The inferred trajectory demonstrated that Grade 1 tumor cells were enriched at the beginning of the trajectory, whereas Grade 2 tumor cells were predominantly located toward the terminal states, suggesting a stepwise progression in chromatin accessibility landscapes (**Fig. 7k**). This spatial trajectory recapitulated the patterns observed in our snATAC-seq dataset (**Fig. 3a**). For instance, FOXA2 expression appeared at the early stage of tumor development in snATAC-seq analysis (**Fig. 3b**), and this trend was similarly observed in the spatial ATAC-seq dataset. Notably, the TF deviation score of FOXA2 was higher in sample PNET1 (Grade 1) compared to sample PNET8 (Grade 2), and the gene activity score of FOXA2 also showed a higher level in PNET1 relative to PNET8 (**Fig. 7m, 7n**), further validating the dynamic transcriptional regulation during NF-PanNET progression.

### myCAF-associated spatial heterogeneity in NF-PanNETs

Previous studies have reported that patients with a high stromal proportion tend to have worse prognosis, particularly when the stroma exhibits an infiltrative growth pattern, where stromal and tumor cells intermix without a clear boundary^65^. Among the stromal components, myCAFs, characterized by high expression of FAP and ACTA2, are known to localize near tumor cells and promote EMT and tumor progression. We identified myCAFs in the snATAC-seq datasets and sought to investigate their spatial relationship with tumor cells using spatial ATAC-seq. After dimensionality reduction and clustering, cell types were annotated based on marker gene expression and pathway enrichment analyses (**Extended Data Fig. 8a-8i**). Interestingly, two distinct tumor subtypes (Tumor1 and Tumor2) were identified in sample PNET8 (**Fig. 8a-8c**). Spatial ATAC-seq analysis revealed that both Tumor1 and Tumor2 were located in close proximity to fibroblasts, with Tumor2 being notably adjacent to regions enriched for high gene activity scores of ACTA2 and FAP, indicating spatial co-localization with myCAFs (**Fig. 8d, 8e**). To quantify this observation, we calculated the spatial distances between myCAFs and the two tumor subtypes, confirming that Tumor2 is spatially closer to myCAFs (**Fig. 8f**). Functional pathway enrichment analysis further revealed that Tumor2 was enriched in proliferative signaling pathways, such as E2F targets, MYC targets, and mTORC1 signaling, alongside oxidative phosphorylation. In contrast, Tumor1 exhibited enrichment of genes involved in the KRAS signaling pathway (**Fig. 8c**).

**Fig. 8.**
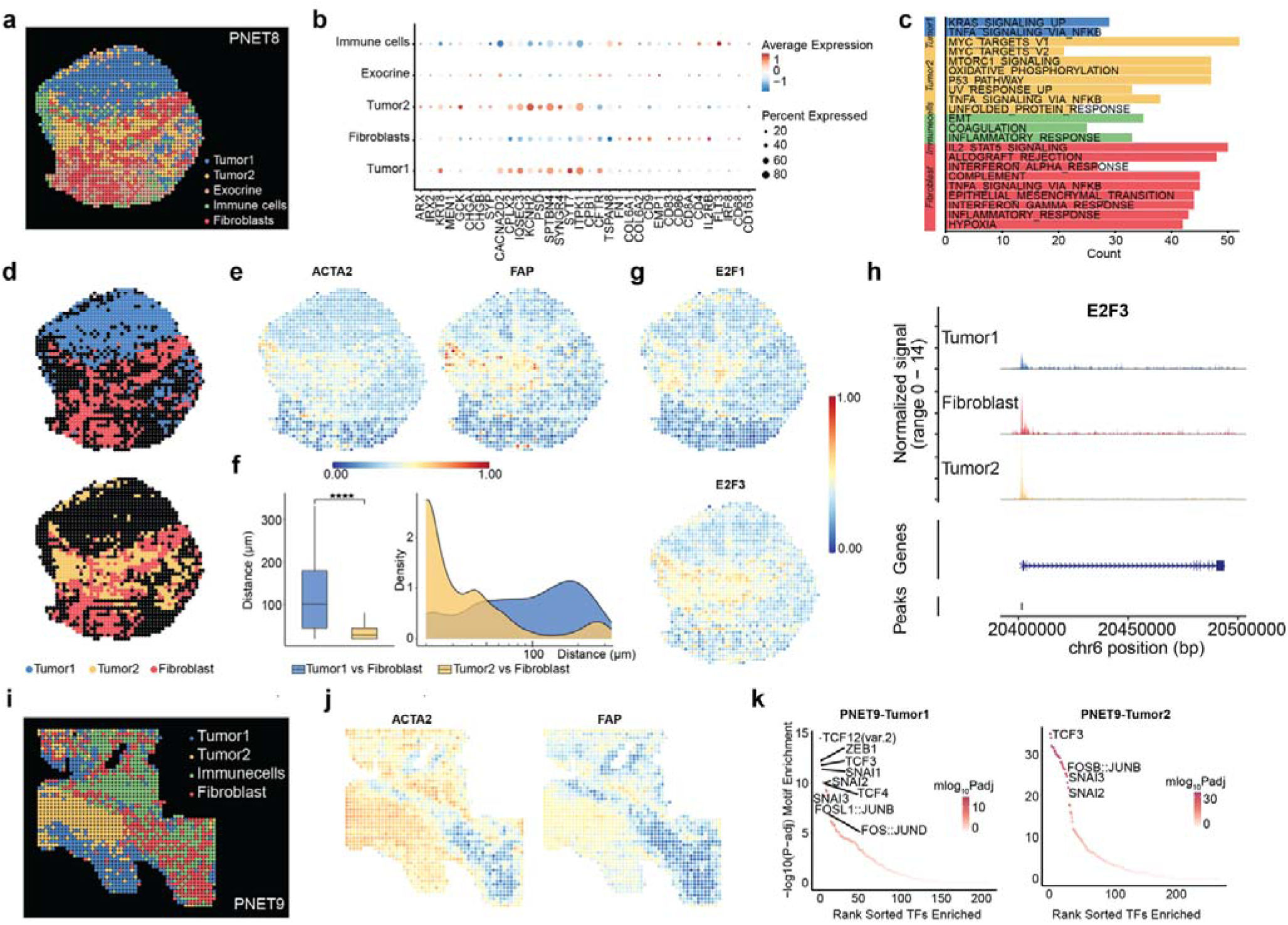
Intra- and intertumoral heterogeneity revealed by spatial chromatin accessibility. a, Spatial distribution of cell clusters in sample PNET8. b, Dot plot showing gene activity scores of representative marker genes across PNET8 subtypes. Dot size indicates the proportion of cells with gene activity; color intensity reflects the average gene activity score. Dark red means higher gene activity score, dark blue means lower gene activity score. c, Enriched hallmark pathways associated with upregulated DACRs in each PNET8 subtype. d, Unsupervised clustering and spatial localization of Tumor1, Tumor2, and Fibroblasts in PNET8. e, Spatial gene activity maps of ACTA2 and FAP in PNET8. f, Distribution of distances between tumor and fibroblast cells in PNET8. Statistical significance was assessed using a two-tailed Mann–Whitney U test (****p < 0.0001). g, Spatial gene activity maps of E2F1 and E2F3 in PNET8. h, Chromatin accessibility profiles at the E2F3 locus across different cell types. i, Spatial distribution of cell clusters in sample PNET9. j, Spatial gene activity maps of ACTA2 and FAP in PNET9. k, Enriched TF motifs in Tumor1 and Tumor2 subclusters of PNET9.

We extended this spatial analysis to other samples (**Extended Data Fig. 10a-10c**). Tumors from PNET1, PNET4, and PNET6 displayed high gene activity scores for ACTA2 and FAP and were enriched in proliferative pathways, similar to Tumor2 in PNET8. In contrast, tumors from PNET7 and PNET10 showed low ACTA2 and FAP activity but were enriched in the KRAS pathway. Notably, intra-tumor heterogeneity was also observed in PNET9, where two distinct tumor subtypes were identified (**Fig. 8i-8j**, **Extended Data Fig. 10d**). As in PNET8, Tumor2 in PNET9 was spatially associated with myCAFs and enriched in proliferative pathways, while Tumor1 was characterized by KRAS pathway activity. In summary, tumors that are spatially proximal to myCAFs tend to activate proliferative transcriptional programs, while those spatially distant from myCAFs are more associated with KRAS signaling. Conversely, myCAFs in proximity to tumors exhibit EMT and hypoxia-related pathway activation.

Further analysis revealed that E2F transcription factors, which regulate cell cycle progression, were upregulated in Tumor2 of PNET8 compared to Tumor1 (**Fig. 8g**). Browser track visualization of E2F3 showed elevated chromatin accessibility in Tumor2 (**Fig. 8h**), supporting its proliferative phenotype. Additionally, motif enrichment analysis across samples showed that FOX motifs were enriched in highly proliferative tumors, consistent with their role in regulating the cell cycle (**Extended Data Fig. 10e**). In contrast, tumors enriched in the KRAS pathway displayed increased occupancy of SNAI1, SNAI2, and SNAI3 motifs—members of the Snail family of zinc-finger transcription factors (**Fig. 8k, Extended Data Fig. 10f-10g**). KRAS activation can induce SNAI1 and SNAI2 expression, which repress epithelial gene expression and promote EMT^66,67^. By binding to their corresponding motifs, these transcription factors orchestrate a transcriptional program downstream of KRAS that supports tumor invasion, metastasis, and apoptosis resistance.

Together, these findings highlight not only intertumor heterogeneity across different patients, but also intratumor heterogeneity within individual tumors, as exemplified in PNET8 and PNET9, where two spatially distinct tumor subtypes exhibit complementary functional programs. This spatial and functional diversity may contribute to tumor growth, immune evasion, and metastasis through collaborative crosstalk between tumor subtypes and the stromal microenvironment. The spatial ATAC-seq data provide a novel framework for identifying high-risk tumor zones within individual tumors. By mapping the spatial distribution of epigenetically active regions, we are able to localize proliferative niches and invasive fronts, each defined by specific regulatory programs (e.g., FOX/MYC versus SNAI/KRAS). Such spatially resolved information has the potential to inform surgical margins, guide biopsy site selection, and enable targeted delivery of therapeutics to the most aggressive tumor compartments.

## DISCUSSION

Our integrative study combining snATAC-seq and spatial ATAC-seq provides an unprecedented, high-resolution epigenomic map of NF-PanNETs. By profiling chromatin accessibility both at single-cell resolution and within the native tissue architecture, we unveil the regulatory mechanisms driving tumor progression and illuminate how these processes are shaped by spatially distinct tumor microenvironments.

Our snATAC-seq analysis reveals that tumor progression in NF-PanNETs is not random but follows an orchestrated, programmatic shift in TF activity. In Grade 1 tumors, chromatin accessibility is enriched for lineage-defining TFs such as HNF1A, HNF4A, and FOXA2, which reflect a well-differentiated, neuroendocrine phenotype. As tumors advance to Grade 2, we observe a striking downregulation of these TFs, concurrent with upregulation of proliferative regulators, including the E2F family and AP-1 complex. This transition parallels a dedifferentiation process common in many solid tumors, but here we provide direct epigenetic evidence specific to NF-PanNETs. Moreover, we observe a sharp increase in YAP1 activity, an effector of the Hippo pathway and a known marker of stemness and poor prognosis in pancreatic cancer, particularly in Grade 3 tumors^68,69^. These findings delineate a stepwise, TF-driven trajectory of NF-PanNET progression - from lineage fidelity in low-grade tumors, through proliferative reprogramming in intermediate grades, to stemness-associated regulatory dominance in high-grade disease. This epigenetic roadmap underscores the potential clinical utility of transcription factor activity profiles as biomarkers of progression, as TF activity such as SNAI and FOXA2 may offer a more accurate molecular indicator of tumor grade and aggressiveness than traditional markers like Ki-67 alone.

Spatial ATAC-seq analysis adds a critical layer of context by linking chromatin states to physical cell locations. This reveals that the spatial positioning of tumor cells relative to stromal elements, particularly CAFs, has profound implications for their epigenetic state and functional behavior. Two distinct tumor–stroma ecological niches were found in NF-PanNETs, proliferative niche and invasive niche. In proliferative niche, tumor cells which are in close proximity to myCAFs, characterized by high FAP and ACTA2 activity, exhibit strong activation of MYC and MTORC1 signaling. These myCAFs display enhanced EMT and hypoxia signaling, suggesting a bidirectional feedback loop that promotes tumor proliferation. In invasive niche, tumor cells spatially segregated from myCAFs exhibit lower proliferative markers but show high activity of Snail family TFs (SNAI1/2/3) and KRAS signaling, indicative of an invasive and potentially migratory phenotype. Notably, such spatial-functional relationships are not limited to inter-tumoral comparisons; we also observe intra-tumor heterogeneity. In samples such as PNET8 and PNET9, distinct tumor subtypes co-exist within the same tumor mass, one subtype closely associated with myCAFs and proliferation, the other more KRAS-driven and spatially separated. These insights emphasize that cellular function in NF-PanNETs is not solely determined by genetic or epigenetic identity, but also by spatial context. We acknowledge certain limitations of the present study, including the relatively small sample size and the limited representation of grade 3 tumors. Neverthelss, our findings offer several promising avenues for clinical translation, particularly in the context of molecular stratification and spatially informed precision oncology. The identification of tumor subtypes defined by distinct TF programs, such as the HNF/FOXA2 axis in well-differentiated tumors versus the E2F/YAP1-driven phenotype in high-grade lesions, establishes a regulatory basis for molecular subtyping beyond conventional histopathology. These TF signatures, inferred from chromatin accessibility profiles, could serve as robust biomarkers for tumor grading, prognosis, and therapeutic response prediction. Moreover, future research should focus on multi-modal data integration, combining chromatin accessibility with spatial transcriptomics, proteomics, and metabolomics to construct a comprehensive, layered view of the tumor ecosystem. This integrative approach will be instrumental in refining tumor classification, uncovering mechanisms of resistance, and identifying novel vulnerabilities. The recurrent activation of TF networks such as the YAP1-HIPPO, FOXA2-HNF, and SNAIL-EMT axes presents compelling opportunities for targeted intervention. Pharmacological inhibition or transcriptional modulation of these regulators may disrupt key epigenetic programs that underlie NF-PanNET progression. Ultimately, this study sets the stage for the development of spatially informed, mechanism-based therapeutic strategies tailored to the specific regulatory architecture and microenvironmental context of each tumor.

## METHODS

### Human participants

All human samples were collected with informed consent in concordance with the Yale University Institutional Review Board (IRB) at the Yale University School of Medicine. Human primary pancreatic neuroendocrine tumor samples were collected during surgical resection and verified by standard pathology (Yale HIC approval number: 1203009817). The tissue collection and optimal cutting temperature (OCT) embedment was conducted with Yale University Institutional Review Board approval with oversight by Tissue Resource Oversight Committee Written informed consent for participation, including cases where identification was collected alongside the specimen, was obtained from patients or their guardians, adhering to the principles of the Declaration of Helsinki. Each sample was handled in strict compliance with HIPAA regulations, University Research Policies, Pathology Department diagnostic requirements, and Hospital by-laws.

### Tissue section preparation and staining

The freshly dissected human samples were immersed into OCT and snapped frozen with liquid nitrogen. Before sectioning, the frozen tissue block was warmed to the temperature of cryotome cryostat (−20□). Tissue block was then sectioned into thickness of 7-10 μm and placed in the center of a poly-L-lysine coated glass slide (electron microscopy sciences, CatLog no. 63478-AS). Serial tissue sections were collected simultaneously for DBiT-ATAC-seq and H&E staining.

The adjacent sections underwent a standard H&E staining procedure. All staining is performed at Yale University School of Medicine, Department of Pathology and at YPTS. These procedures adhered to Clinical Laboratory Improvement Amendments (CLIA)-certified laboratory protocols as well as YPTS’s rigorous standard protocols, ensuring precision and accuracy in the analysis of tissue samples.

### Single-nucleus ATAC (snATAC) assay

OCT-embedded tissues were sectioned to generated 5-10 50um scrolls using a microtome (Finesse ME+, Thermo Scientific, UK). Single cell suspensions were generated from the scrolls using10X Chromium Nuclei Isolation Kit (1000493, 10x Genomics, US) according to the manufacturer’s recommendations. Nucleus suspensions were then flow-sorted with the BD FACSMelody, where 7AAD intensity was used to gate the desired populations.

The snATAC-seq libraries were prepared using the Chromium Next GEM Single Cell ATAC Reagent kits (1000390, 10x Genomics, US) according to the standard protocol. Initially, the isolated nucleus suspensions were incubated in a transposition mix containing a transposase enzyme, facilitating preferential fragmentation of DNA in open chromatin regions and introducing adapter sequences. Subsequently, approximately 10000 transposed nuclei were loaded onto the Chromium Next Chip H. During GEM generation, gel beads introduced a spacer sequence facilitating barcode attachment to transposed DNA fragments allowing individual nuclei to be uniquely labeled during subsequent amplification. After GEM incubation, DNA fragments were purified following the standard protocol, and the purified DNA fragments were amplified via PCR to generate sufficient material for sequencing. Following quality assessment, the snATAC library underwent paired-end 150bp read sequencing on the Illumina NovaSeq 6000 sequencing system.

### Spatial ATAC-seq

DNA oligos including barcodes, barcodes linkers, indexes and transposome oligos are all procured from integrated DNA Technologies (IDT, Coralvile, IA) and the sequences were listed in **Extended Data Table 1-3**. The transposome was assembled beforehand following the manufacturer’s guidelines using unloaded Tn5 transposase (Diagenode, no. C01070020). All other key reagents used were listed in **Extended Data Table 4**. OCT-embedded tissue sections were taken from the −80°C freezer and allowed to return to room temperature for 5-10 minutes. The sections were then fixed with 0.2% formaldehyde in DPBS for 5 minutes and quenched with 1.25 M glycine for an additional 5 minutes. After fixation, the tissue was washed twice with 1 ml of DPBS, and the fixed tissues were then permeabilized for 20 minutes at room temperature with freshly made permeabilization buffer (10 mM Tris-HCl, pH 7.4, 10 mM NaCl, 3 mM MgCl2, 0.01% Tween-20, 0.01% NP-40, 0.001% digitonin, 1% BSA). 1ml wash buffer (10 mM Tris-HCl, pH 7.4, 10 mM NaCl, 3 mM MgCl2, 0.01% Tween-20, 1% BSA) was used to wash the tissue twice after permeabilization. The tissues were then quickly rinsed with deionized water, air-dried and installed with PDMS reservoir and clamps for tagmentation. For each sample, 100ul transposition mix containing 50 µl 2× tagmentation buffer, 33 µl 1× DPBS, 1 µl 10% Tween-20, 1 µl 1% digitonin, 5 µl transposome,10 µl nuclease-free H2O was added followed by incubation at 37 °C for 30 min. After tagmenetation, 100ul 40mM EDTA was added to the tissues instantly to stop the transposition. Finally, the EDTA was removed, and tissues were washed with DPBS for three times, rinsed with water and air-dried for imaing. Spatial barcodes are made days before the tissue processing. Barcode (100 μM) and ligation linker (100 μM) were annealed at a 1:1 ratio in 2X annealing buffer (20 mM Tris-HCl pH 8.0, 100 mM NaCl, 2 mM EDTA) with the following PCR program: 95°C for 5 min, slow cooling to 20°C at a rate of −0.1°C/s, followed by 12°C for 3 min. Barcode ligation was performed through three consecutive procedures: whole image capture, first PDMS device attachment, imaging, loading barcode A (A1-A50) and second PDMS device attachment, imaging, loading barcode B (B1-B50). For downstream alignment and analysis, tissues were scanned three times using a 10X objective on the EVOS M7000 Imaging System. The first image was taken before placing the first PDMS device, covering the entire tissue area, while the second and third images, covering only the ROI, were taken after attaching the first and second PDMS devices, respectively. Microfluidic barcoding was performed by positioning the PDMS device channels over the ROI, with the PDMS device and tissue slide clamped tightly using a homemade acrylic clamp. The ligation mix for barcodes A and B was prepared with 69.5 μL of RNase-free water, 27 μL of T4 DNA ligase buffer (10X, New England Biolabs), 11 μL T4 DNA ligase (400 U/μL, New England Biolabs), and 5.4 μL of Triton X-100 (5%). A 5 μL ligation mix containing 4 μL of the ligation mix and 1 μL of 25 μM DNA barcode A (A1-A50) or barcode B (B1-B50) was added to each of the 50 channels’ inlets and pulled in using a house vacuum for less than 3 minutes. The reaction occurred at room temperature for 30 minutes in a wet box. After reaction, the clamp was removed, the PDMS was carefully peeled off, and the slide was washed for 5 minutes in a Falcon tube containing 50 mL DPBS and air-dried for the next barcoding or lysis. After the second barcoding, a square well PDMS gasket was placed over the ROI where both barcodes had flowed. The reservoir created by the PDMS gasket was loaded with 50-100 μL of freshly made lysis buffer (0.4 mg/ml proteinase K, 1 mM EDTA, 50 mM Tris-HCl pH 8.0, 200 mM NaCl, 1% SDS), depending on the reservoir size. The slides were incubated in a wet box at 55°C for 2 hours. The lysates in the reservoir were then collected in a 1.5 mL DNA low-binding tube and incubated at 55°C overnight. Genomic fragments in the lysate were purified using the Zymo DNA Clean & Concentrator-5 and eluted into 22 μL of RNase-free water. The ATAC library was built following a direct PCR. The PCR mix, consisting of 20 μL DNA solution, 25 μL of 2X NEBNext Master Mix, 2.5 μL of 25 μM indexed i7 primer, 2.5 μL of 25 μM P5 PCR primer, and 2.5 μL 20X EvaGreen, was run with the following program in a qPCR machine: 72°C for 5 min, 98°C for 30 s, followed by cycles of 98°C for 10 s, 63°C for 30 s, and 72°C for 1 min. The reaction was stopped when the signal began to plateau. The PCR product was purified using SPRIselect beads at a 1:1 ratio and eluted into 20 μL of nuclease-free water. Finally, the library quality was checked using Agilent Bioanalyzer High Sensitivity Chips, and next-generation sequencing (NGS) was conducted on an Illumina NovaSeq 6000 sequencer (paired-end, 150-base-pair mode).

### snATAC data preprocessing

Single-nucleus ATAC-seq (snATAC-seq) data were processed using Cell Ranger ATAC (v2.0, 10x Genomics, https://support.10xgenomics.com), including read alignment, removal of PCR duplicates, peak calling, and generation of fragment counts in a cell-by-peak matrix for each sample. For downstream chromatin accessibility analysis, we used the filtered peak_bc_matrix.h5, singlecell.csv, and fragments.tsv files as input to the Signac package (v1.12.0) in R^70^. Low-quality cells with a transcription start site (TSS) enrichment score < 3 or fewer than 500 unique nuclear fragments were excluded from further analysis.

### Spatial ATAC-seq data preprocessing

Spatial ATAC-seq data were preprocessed using a previously published pipeline (https://github.com/dyxmvp/Spatial_ATAC-seq). Briefly, Read 1 was filtered using Linker 1 and Linker 2. The filtered sequences were then reformatted into the Cell Ranger ATAC (10x Genomics) input format. In the new format, genomic sequences were retained in Read 1, while Barcode A and Barcode B were incorporated into Read 2. The resulting FASTQ files were aligned to the human reference genome (GRCh38), followed by duplicate removal and fragment counting using Cell Ranger ATAC v2.0. Fragment files were generated for downstream analysis, with each entry containing genomic coordinates and the corresponding spatial barcode information (Barcode A × Barcode B).

### Integrative ATAC-seq data analysis

Integrated analysis of snATAC-seq/spatial ATAC-seq datasets was performed using Signac (v1.12.0) and Seurat (v5.0.1). Chromatin accessibility profiles were merged using the Merge function in Signac. Prior to merging, a unified peak set was constructed by aggregating peaks from all samples, followed by quantification in each dataset. Feature selection was performed using the FindTopFeatures function. To correct for batch effects across samples, the Harmony algorithm was applied with default parameters. Data were normalized and dimensionally reduced using latent semantic indexing (LSI), followed by graph-based clustering and UMAP embedding using the FindNeighbors, FindClusters, and RunUMAP functions in Seurat. UMAP visualizations were generated using the DimPlot function.

### Gene activity and TF dynamics analysis

Gene activity scores were calculated using Cicero^30^, which aggregates accessibility signals across gene promoters and their cis-co-accessible distal regulatory elements. TF activity was assessed using the RunChromVAR function in Signac, based on position weight matrices (PWMs) from the chromVARmotifs R package^32^. Per-cell motif deviation scores were used to infer relative TF activity across cell types. Differentially accessible chromatin regions (DACRs) and TF motifs were identified using the FindAllMarkers function (min.pct = 0.1, logfc.threshold = 0.25).

### Cell cluster annotation

Cell clusters were annotated by comparing gene activity scores of known marker genes. Tumor cells were defined by expression of ARX, IRX2, KRT8, MEN1, GCK, CHGA, CHGB, SYP, CACNA2D2, CPLX2, IQSEC3, KCNH2, PSD, SPTBN4, SYNGR4, SYT7, and ITPK1. Exocrine cells were marked by CPB1, CFTR, and TSPAN8. Fibroblasts were identified via FN1, COL6A1, COL6A2, CD9, and EMP1. Immune cell types were annotated as follows: B cells (CD83, CD86, CD19, MS4A1, CD79A, CD79B), T cells (CD8A, CD8B, CD4), NK cells (IL2RB, SPON2, KLRD1, CD160), conventional dendritic cells (cDCs; FLT3, IRF8), and macrophages (CD68, CD163, LAMP2, ITGAM, LGALS3). CAF subtypes were defined as follows: inflammatory CAFs (iCAFs; IL6, CXCL12), myofibroblastic CAFs (myCAFs; FAP, ACTA2), and antigen-presenting CAFs (apCAFs; CD74, HLA-DPB1, HLA-DRA, HLA-DRB1).

### Identification and annotation of cell-type specific chromatin accessible regions

Cell type–specific accessible chromatin regions (peaks) were identified using the FindAllMarkers function in Seurat, with the following thresholds: log fold change > 0.5, minimum of 50 cells per group, and adjusted *P*[<[0.05. Peaks were annotated to the nearest genes using the annotatePeak function in the ChIPseeker R package, with promoter regions defined as ±3 kb from the transcription start site (TSS). Chromatin states were annotated using the core 15-state model from the NIH Roadmap Epigenomics Mapping Consortium (https://egg2.wustl.edu/roadmap/web_portal/chr_state_learning.html#core_15state). ChromHMM states were assigned to peak regions using the annotatedPeak function from ChIPpeakAnno with default parameters.

### Motif enrichment analysis

Motif enrichment was performed using a hypergeometric test implemented in the FindMotifs function of Signac, comparing the frequency of TF motifs in a target peak set versus background peaks matched for GC content.

### TF footprinting analysis

TF footprinting analysis was conducted using Signac. Motif positions were first added to the peak matrix. The Footprint function was used to compute the normalized observed/expected Tn5 insertion frequency surrounding each motif instance across the genome. The resulting footprints were visualized using the PlotFootprint function.

### Pseudotime analysis

To assess gene and TF activation dynamics along tumor differentiation, we constructed pseudotime trajectories and evaluated their significance based on cluster ordering, as previously described^71^. Gene and TF activity scores were smoothed along pseudotime using the loess function in R (span = 2). Heatmaps and trajectory plots were generated using the ggplot2 package in R to visualize the dynamic changes in gene and TF activities.

### Genomic track profile

Chromatin accessibility of specific genomic regions was visualized by function CoveragePlot in Signac.

### Pathway enrichment analysis

Pathway enrichment analysis was performed by first setting the default assay to “RNA” using the DefaultAssay function. Differentially expressed genes (DEGs) for each cluster were identified using the FindAllMarkers function with a minimum expression threshold of 10% (min.pct = 0.1) and a log fold change cutoff of 0.25. Gene set enrichment was conducted using the enricher function from the clusterProfiler^72^ package against the 50 hallmark gene sets curated in the Molecular Signatures Database (MSigDB)^73^.

### Spatial ATAC data visualization

Spatial pixel positions were initially determined from brightfield images using MATLAB 2020b (https://github.com/edicliuyang/Hip-lex_proteome). To visualize spatial distributions of chromatin features (e.g., unique nuclear fragments), fragment files were processed with ArchR^74^ (v1.0.1) to generate an ArchRProject, which was then imported into Seurat (v5.0.1) for tissue-level mapping. The pt.size.factor parameter was used to scale pixel size for optimal visualization. For gene activity scores or TF deviation scores, metadata from Signac objects were extracted and spatially projected onto tissue sections accordingly.

### Statistical analysis

Statistical analyses were performed using Prism10. Differences between two groups were assessed using unpaired, two-tailed Student’s *t*-tests. *P* values less than 0.05 were considered statistically significant.

## RESOURCE AVAILABILITY

## Data availability

The snATAC and spatial ATAC sequencing data generated in this study have been deposited in public repositories to ensure transparency and reproducibility. They are available through the NCBI Gene Expression Omnibus (GEO) under accession number GSE304813.

## Code availability

All custom scripts used for data preprocessing, analysis, and visualization are freely accessible at the following GitHub repository: https://github.com/Xiangjun99/NF-PanNETs-ATAC-project.

## Acknowledgements

The authors acknowledge the support from Department of Pathology and Department of Neurology at Yale School of Medicine. We also thank the Yale Pathology Tissue Services. The research is supported by Maximizing Investigators Research Award (MIRA) for Early Stage Investigators R35 GM150838 and NIH R01 HL173271 (to Y.L.). The research is also supported, in part, by Grant #IRG-21-132-60-IRG from the American Cancer Society. We also acknowledge the support of Leslie H. Warner Postdoctoral Fellowship to D.W.

## Author contributions

Conceptualization: Y.L., J.D., D.W. Methodology: D.W., X.D., Y.L., F.G., G.L., J.D., Y.J. Investigation: D.W., X.D., Y.L. Visualization: X.D., Y.L. Writing – original draft: X.D., D.W. Writing – review & editing: X.D., D.W., Y.L., J.D. Resources: F.G., G.L., L.L., S.H., D.Z., J.X., Y.J., J.L., P.K., J.K. Supervision: J.D., Y.L. Funding acquisition: J.D., Y.L.

