## Supplemental information for "Epigenomic Heterogeneity of Non-Functional Pancreatic Neuroendocrine Tumors Uncovered by Single nucleus and Spatial ATAC Profiling"

^7^Leader contact

**Table of Contents**

**3** Extended Data Figs. 1-10

**13** Extended Data Table 1-4

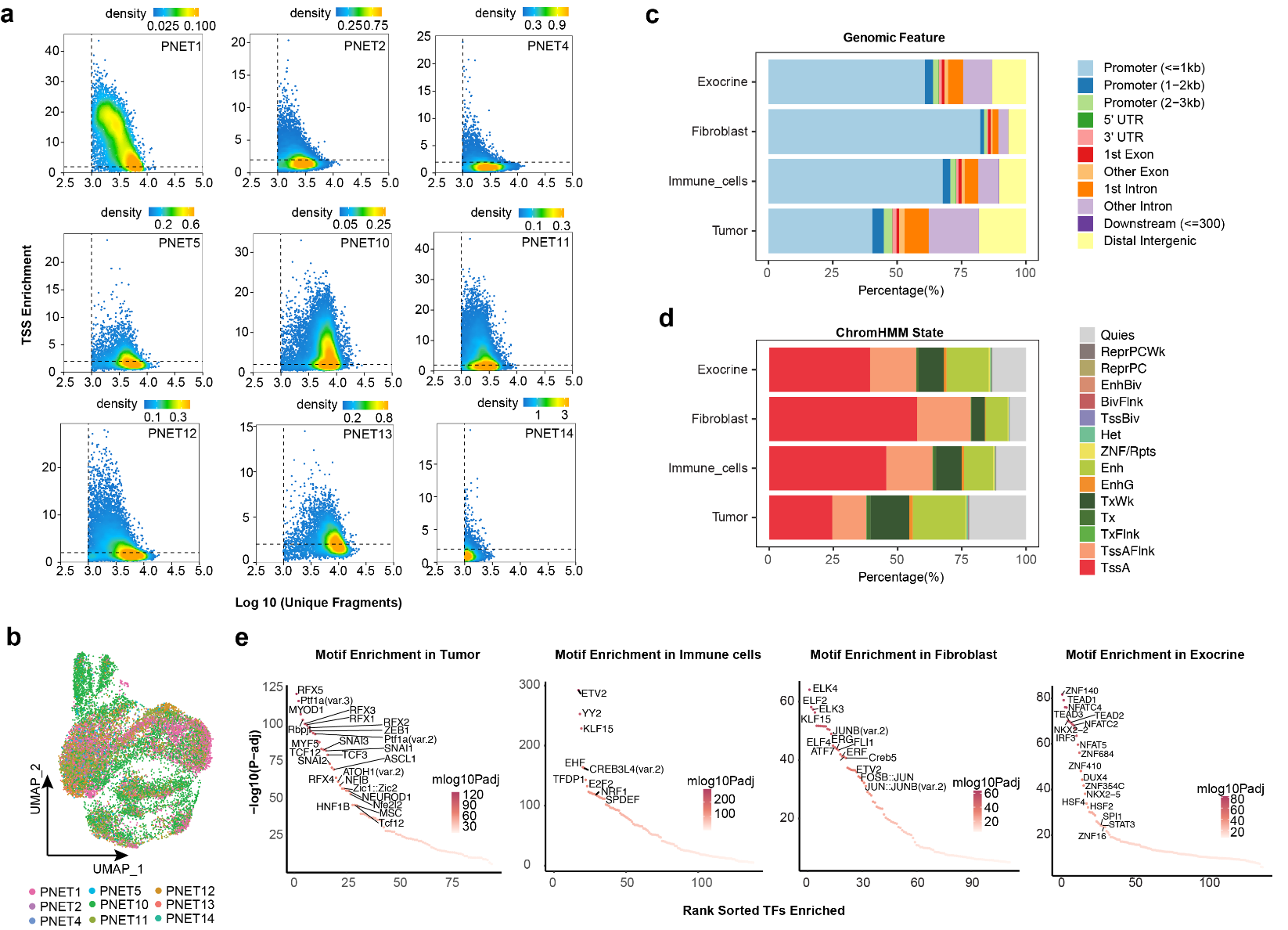

**Extended Data Fig. 1 Extended analysis of snATAC-seq profiling across nine NF-PanNET samples.** a, Scatter plot of TSS enrichment score versus unique nuclear fragments per cell across nine NF-PanNETs, reflecting data quality and chromatin accessibility. b, UMAP of snATAC-seq profiles across samples. c, Genomic feature distribution of differentially accessible chromatin regions (DACRs) specific to each cell type. d, ChromHMM-based chromatin state annotations for cell type-specific DACRs, categorized as follows: TssA (Active TSS), TssAFlnk (Flanking Active TSS), TxFlnk (Transcription at gene 5′ and 3′ ends), Tx (Strong transcription), TxWk (Weak transcription), EnhG (Genic enhancers), Enh (Enhancers), ZNF/Rpts (ZNF genes and repeats), Het (Heterochromatin), TssBiv (Bivalent/Poised TSS), BivFlnk (Flanking Bivalent TSS/Enhancer), EnhBiv (Bivalent Enhancer), ReprPC (Repressed PolyComb), ReprPCWk (Weak Repressed PolyComb), and Quies (Quiescent/Low signal). e, Enriched transcription factor (TF) motifs identified in cell type-specific DACRs, suggesting distinct regulatory programs across cell populations.

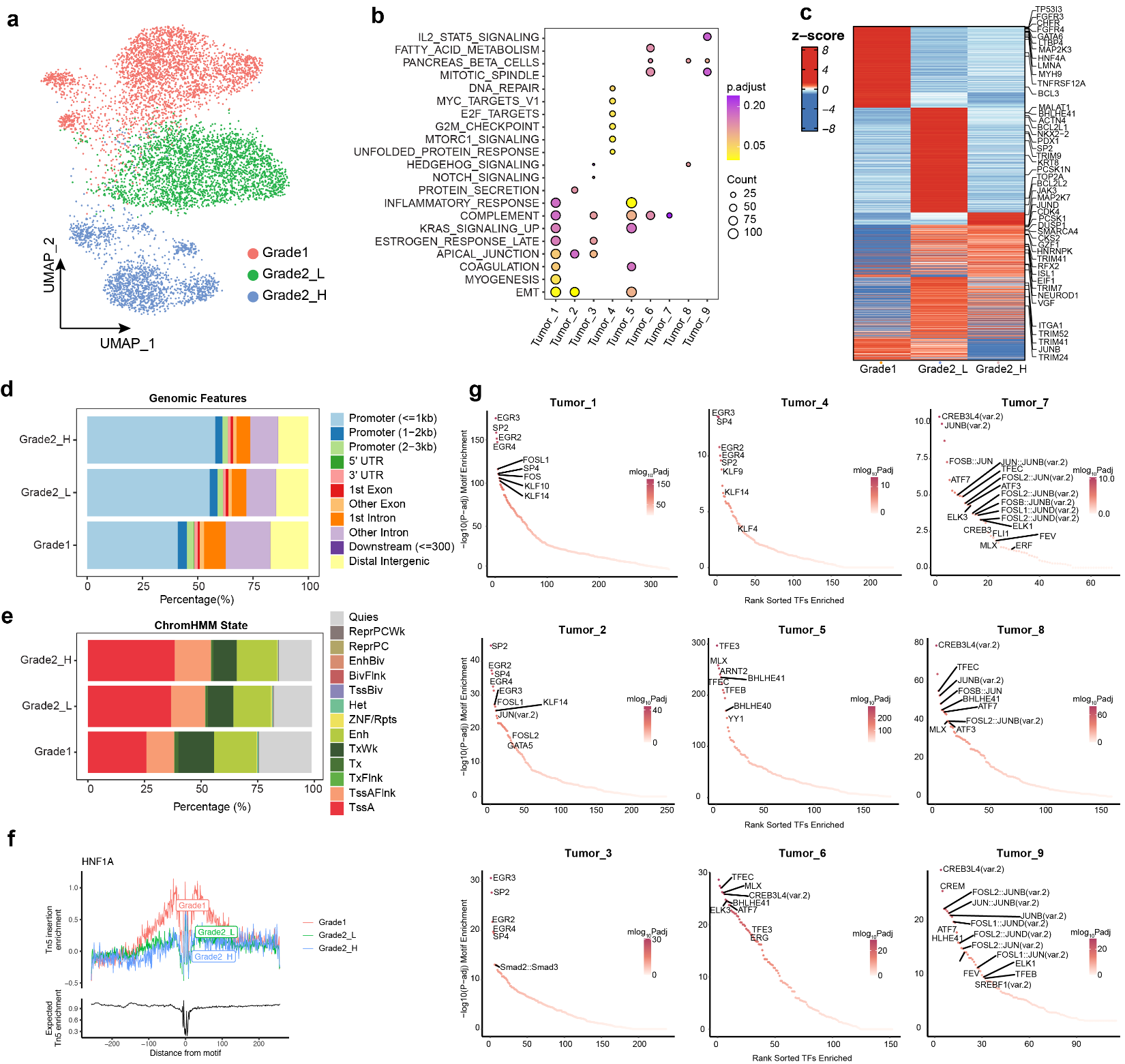

**Extended Data Fig. 2. Further analysis of chromatin accessibility and TF activities in tumor cells across grades.** a, UMAP showing the annotated spatial-ATAC-seq profiles of tumor cells from different grades. b, Enrichment of hallmark pathways derived from DACRs upregulated in each tumor subtype. Bubble size indicates gene count, and color represents the adjusted p-value. c, Top differentially accessible peaks (DAPs) identified in tumor cells for each grade. Color indicates z-scaled average normalized accessibility. d, Genomic feature distribution of tumor grade-specific DACRs. e, ChromHMM annotation of tumor grade-specific DACRs. TssA, Active TSS; TssAFlnk, Flanking Active TSS; TxFlnk, Transcription at gene 5′ and 3′; Tx, Strong transcription; TxWk, Weak transcription; EnhG, Genic enhancers; Enh, Enhancers; ZNF/Rpts, ZNF genes and repeats; Het, Heterochromatin; TssBiv, Bivalent/Poised TSS; BivFlnk, Flanking Bivalent TSS/Enhancer; EnhBiv, Bivalent Enhancer; ReprPC, Repressed PolyComb; ReprPCWk, Weak Repressed PolyComb; Quies, Quiescent/Low. f, HNF1A footprinting profiles across tumor grades. g, Motif enrichment analysis of subtype-specific DACRs.

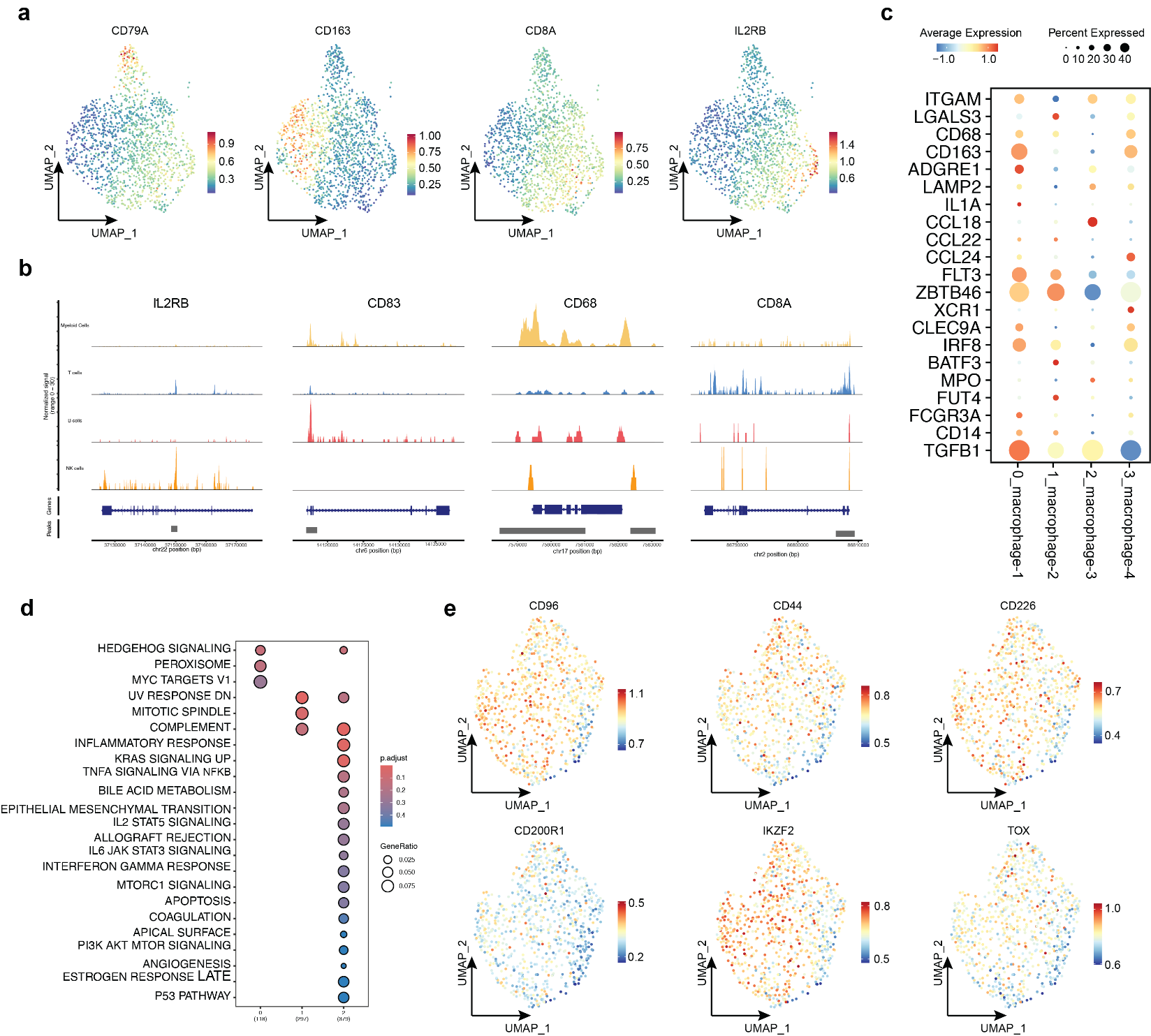

**Extended Data Fig. 3 Further analysis of chromatin accessibility and TF activities in immune cells.** a, UMAP of gene activity scores for selected marker genes across immune cell subtypes. b, Coverage plots showing chromatin accessibility at marker gene loci for each immune cell subtype. c, Dot plot of marker gene activity scores in macrophage subtypes. Dot size indicates the percentage of cells expressing the gene; color intensity reflects average gene activity scores (dark red: high, dark blue: low). d, Enriched hallmark pathways derived from DACRs upregulated in each T cell subtype. Bubble size and color represent gene ratio and adjusted *p*-value, respectively. e, UMAP projection of gene activity scores for selected marker genes across T cell subtypes.

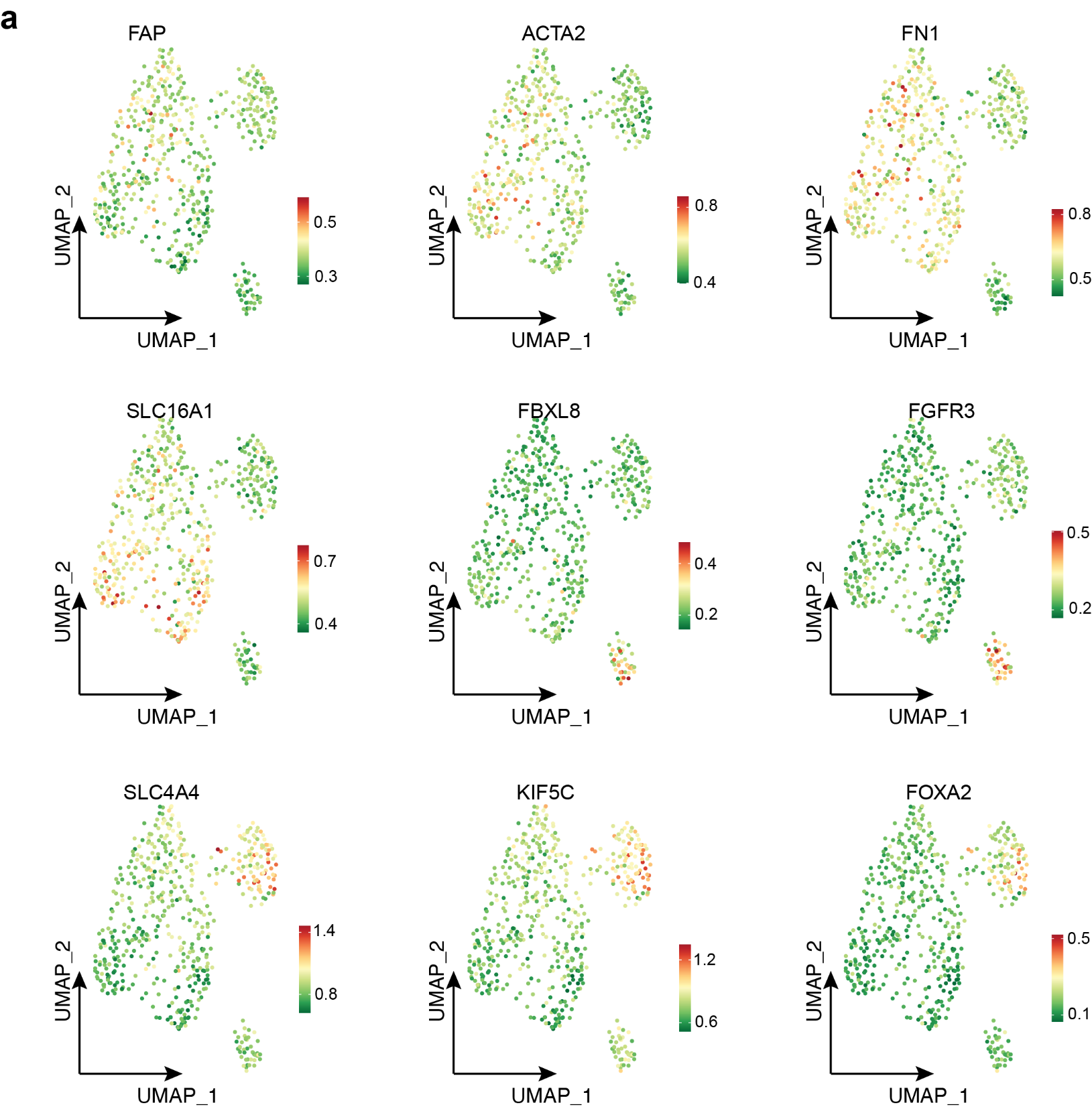

**Extended Data Fig. 4 Further analysis of chromatin accessibility and TF activities of CAFs.** UMAP projection of gene activity scores for selected marker genes across different CAF subtypes.

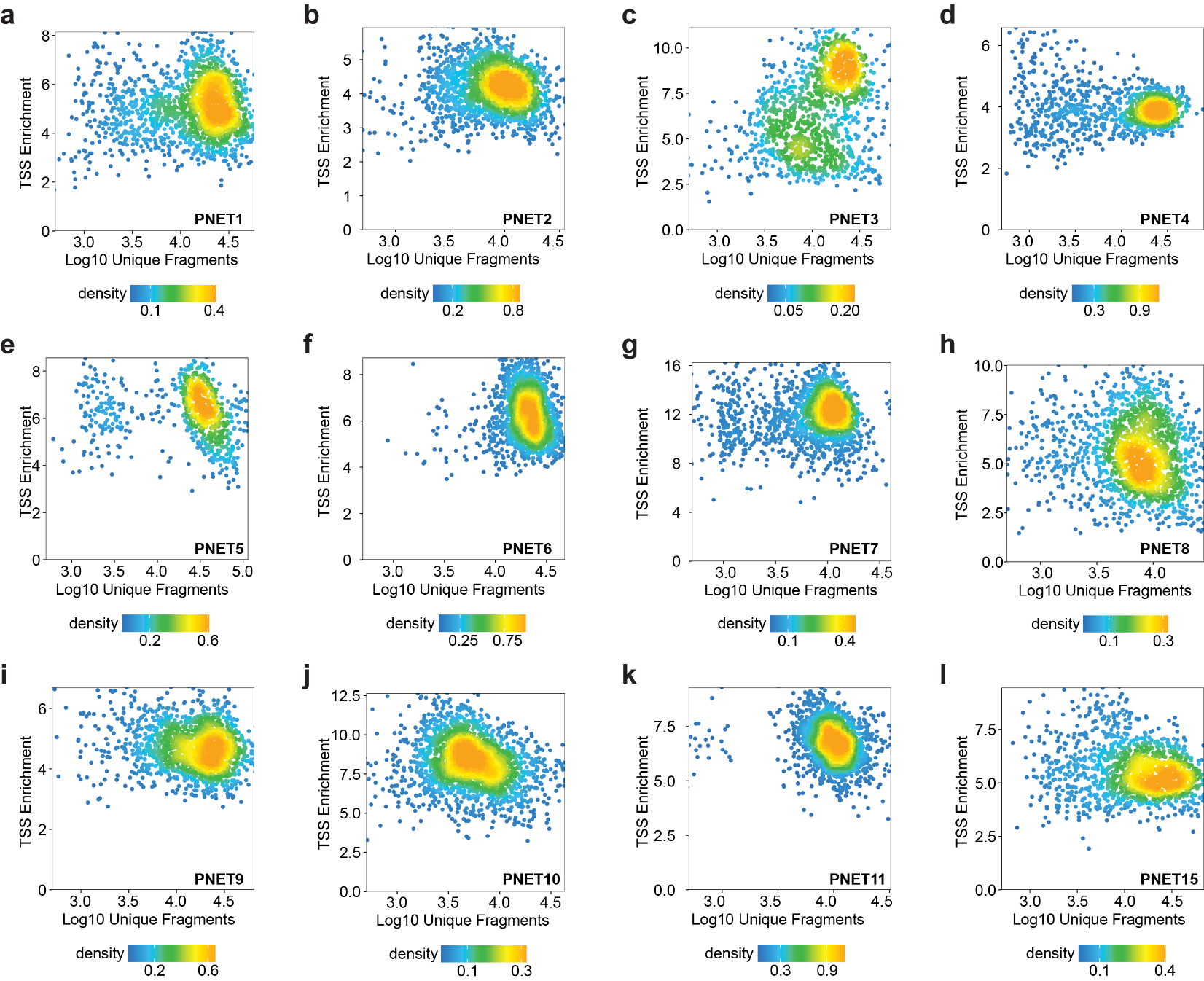

**Extended Data Fig. 5 TSS enrichment score versus unique nuclear fragments per cell for each spatial-ATAC-seq sample.** a, PNET1; b, PNET2; c, PNET3; d, PNET4; e, PNET5; f, PNET6; g, PNET7; h, PNET8; i, PNET9; j, PNET10; k, PNET11; l, PNET15.

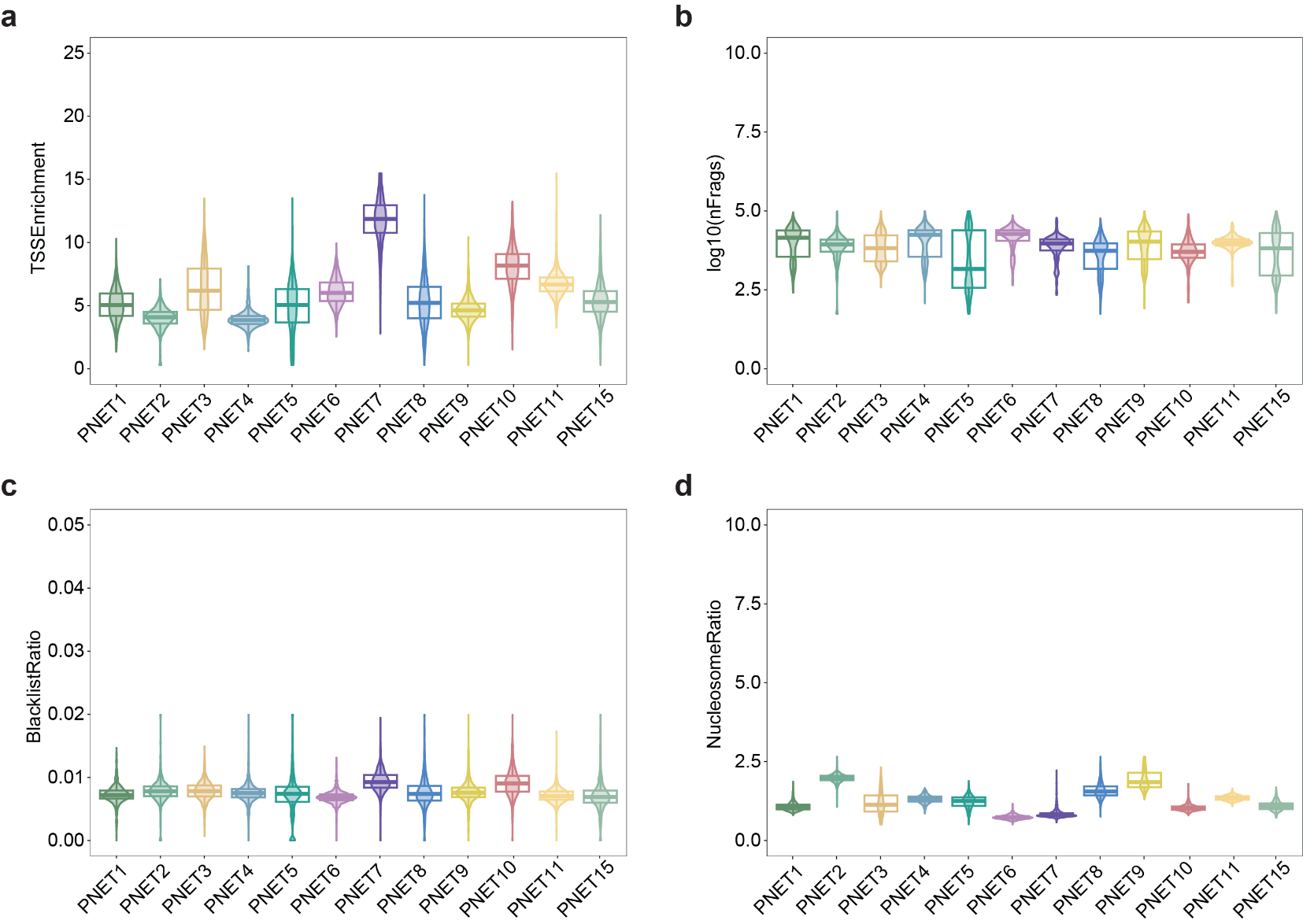

**Extended Data Fig. 6 Data quality metrics of spatial-ATAC-seq across different samples.** a, TSS enrichment scores; b, Unique nuclear fragment counts; c, Blacklist ratios; d, Nucleosome signal ratios.

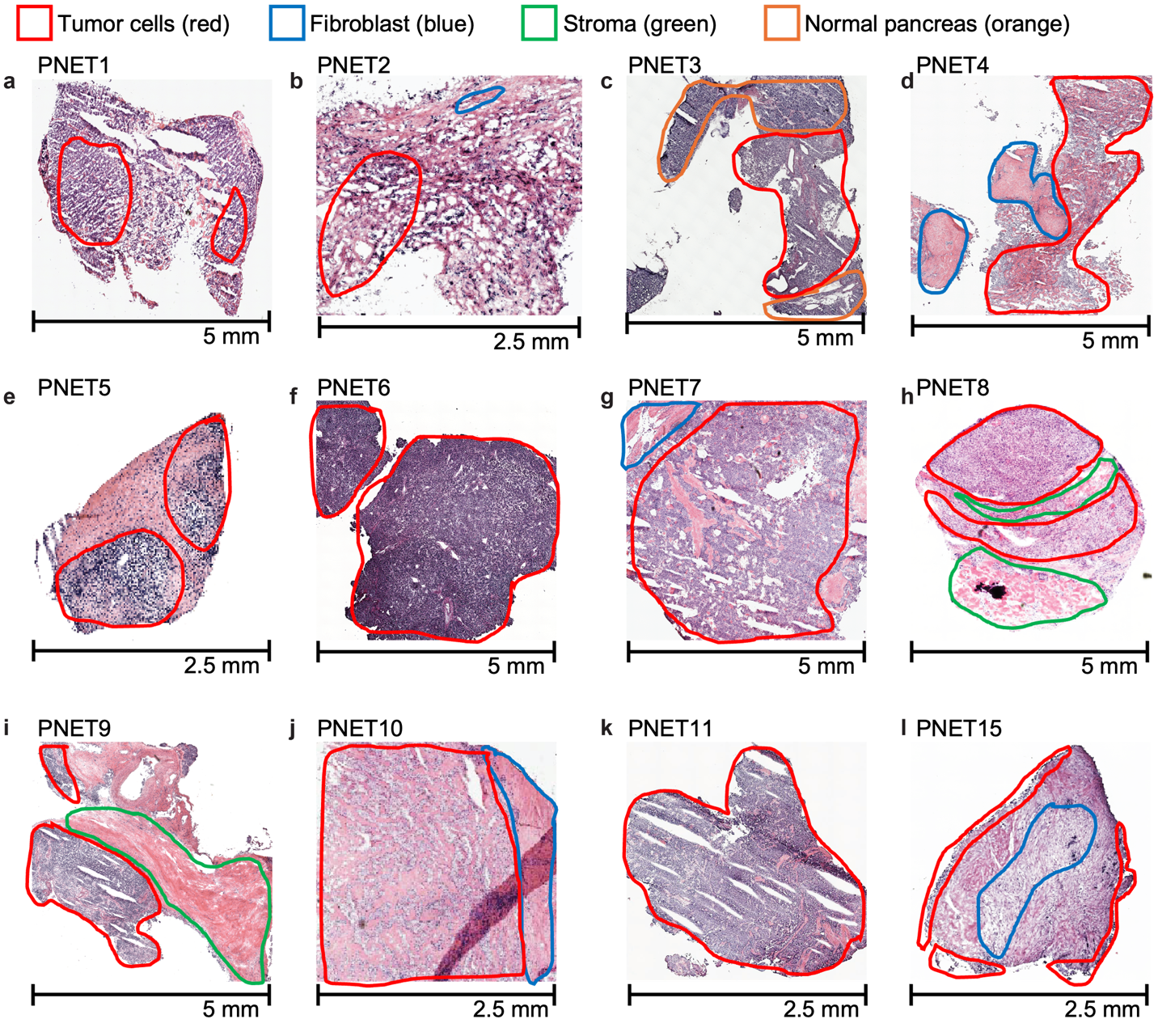

**Extended Data Fig. 7 Evaluating the tissue context of NF-PanNETs using spatial-ATAC-seq.** a–i, H&E-stained images of tumor sections processed for spatial-ATAC-seq. Pathologist-annotated regions are highlighted: tumor (red), fibroblasts (blue), stroma (green), and normal pancreas (orange).

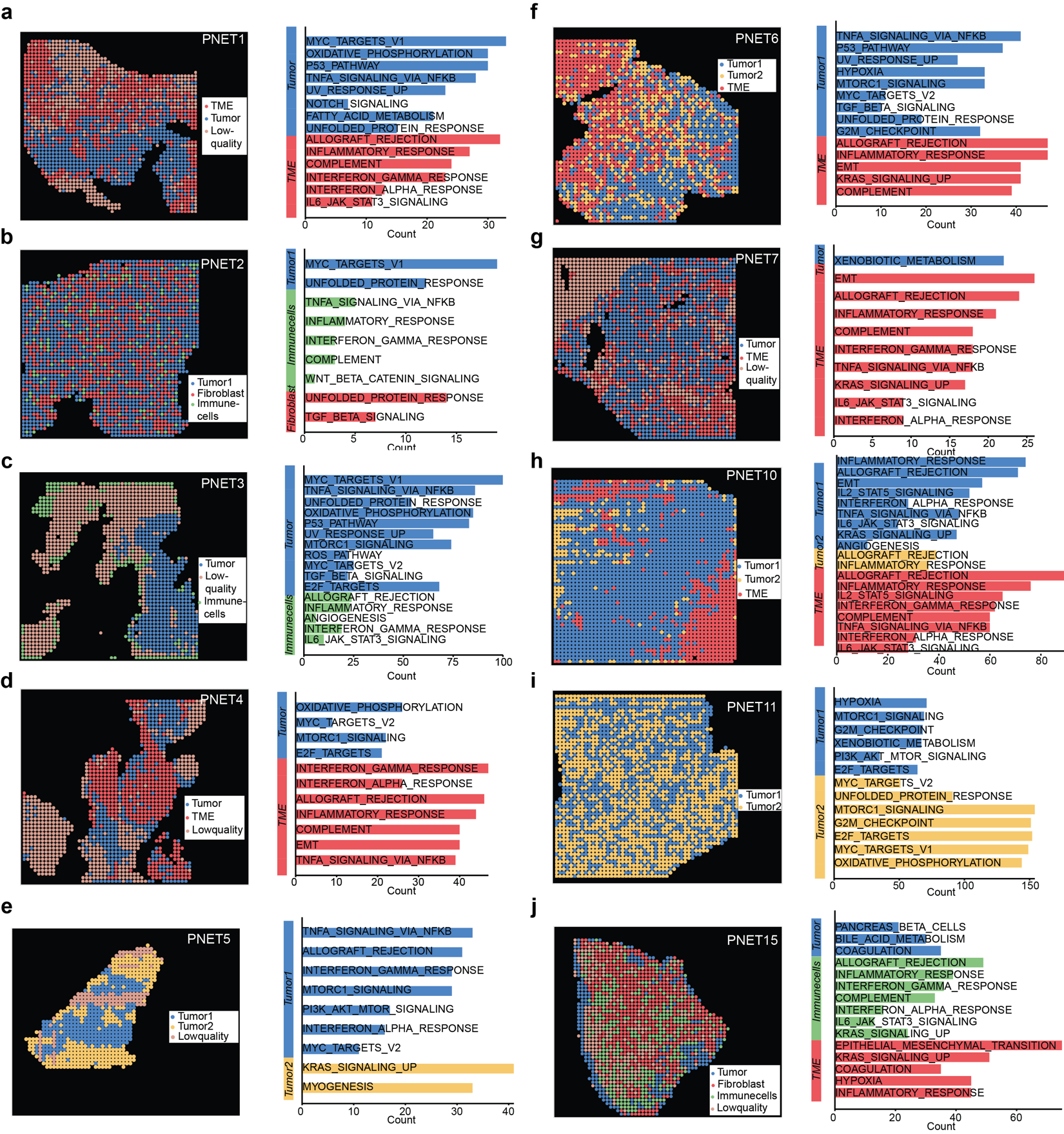

**Extended Data Fig. 8 Spatial mapping of chromatin accessibility-based clusters across NF-PanNET samples.** Unbiased clustering was performed based on chromatin accessibility profiles in each sample: PNET1 (a), PNET2 (b), PNET3 (c), PNET4 (d), PNET5 (e), PNET6 (f), PNET7 (g), PNET10 (h), PNET11 (i), and PNET15 (j).

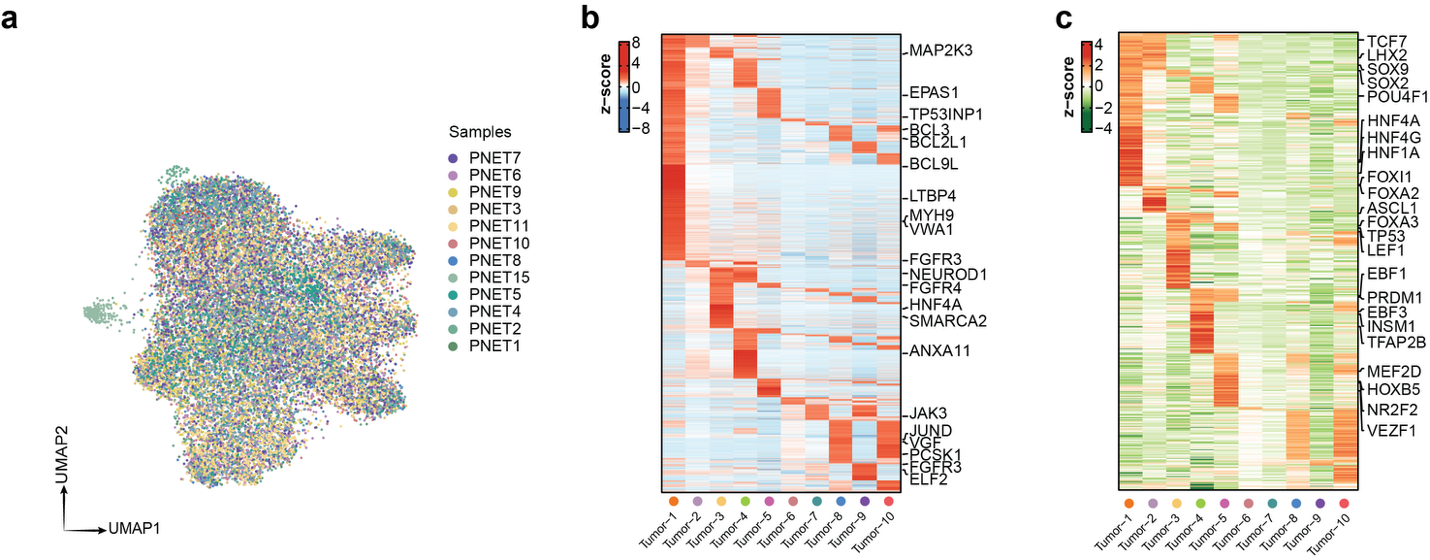

**Extended Data Fig. 9 Spatial chromatin accessibility and regulatory landscape of tumor subtypes in NF-PanNET.** **a,** UMAP projection showing the annotated spatial-ATAC-seq profiles across different samples. **b,** Top DACRs identified for each tumor subtype. Color represents the z-scaled average normalized chromatin accessibility signal within each subtype. **c,** Differentially activated TFs across tumor subtypes. Color indicates the z-scaled average TF activity score for each subtype.

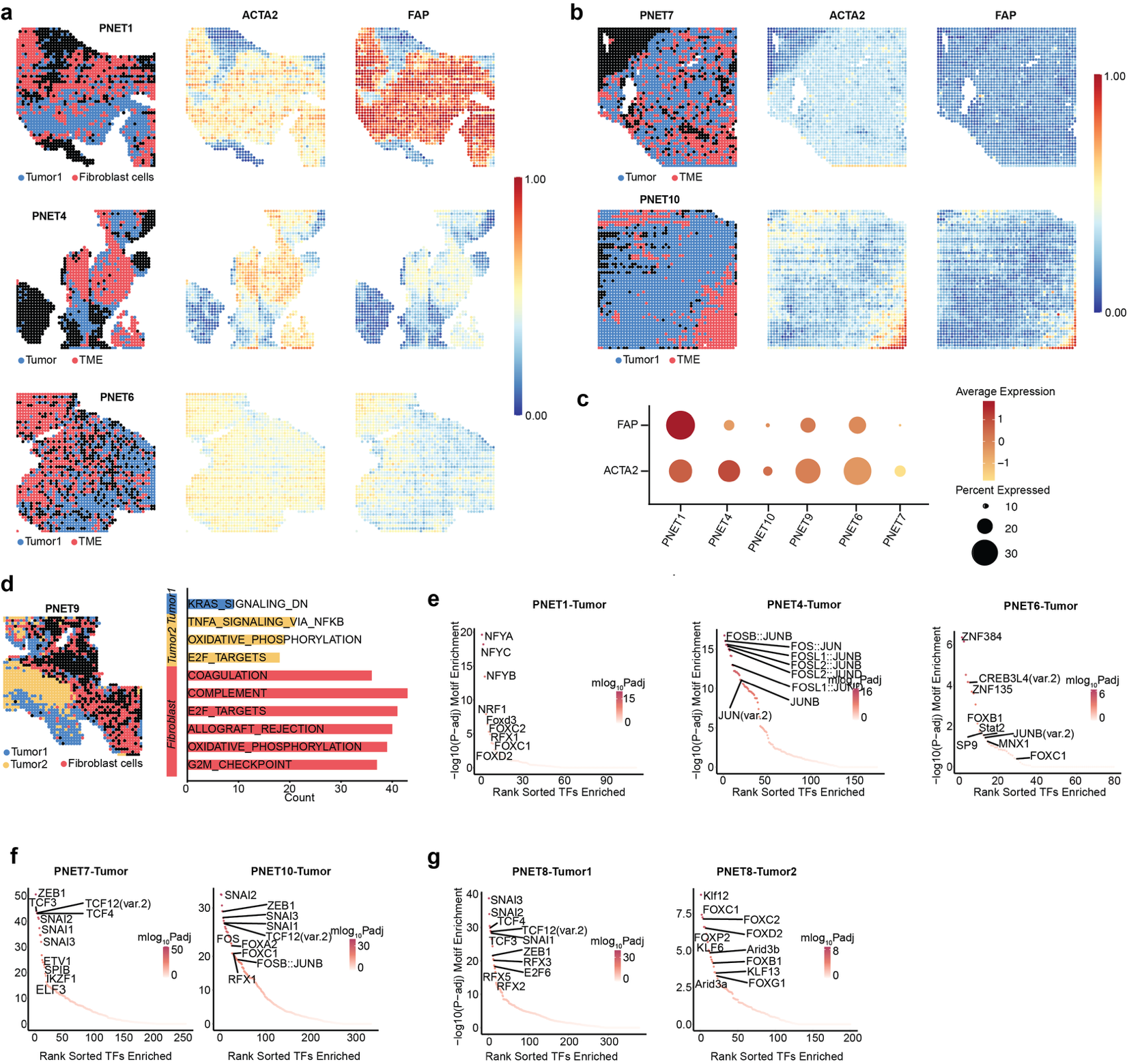

**Extended Data Fig. 10 Further analysis of spatial distribution of chromatin accessibility landscape in stroma and tumor.** a–b, Spatial mapping of tumor and fibroblasts clusters (left), gene activity score of ACTA2 (middle), and gene activity score of FAP (right). c, Dot plot showing the gene activity scores of CAF marker genes across samples. Dot size represents the percentage of cells with detected expression; color intensity indicates the average gene activity score per sample (dark red: higher activity; light red: lower activity). d, Spatial mapping of tumor and fibroblasts clusters (left) and corresponding pathway enrichment analysis (right) in PNET9. e, Enriched TF motifs in tumor-specific DACRs from samples PNET1 (left), PNET4 (middle), and PNET6 (right). f, Enriched TF motifs in tumor-specific DACRs from samples PNET7 (left) and PNET10 (right). g, Enriched TF motifs in tumor-specific DACRs from sample PNET8.

**Extended Data Table 1. DNA oligos used for PCR and preparation of sequencing library.**

| **Sequence Name** | **Sequence (5'-3')** |
| --- | --- |
| Ligation linker 1 | AGTCGTACGCCGATGCGAAACATCGGCCAC |
| Ligation linker 2 | CGAATGCTCTGGCCTCTCAAGCACGTGGAT |
| Tn5MErev | /5Phos/CTGTCTCTTATACACATCT |
| Tn5ME-A | TCGTCGGCAGCGTCAGATGTGTATAAGAGACAG |
| Tn5ME-B | /5Phos/CATCGGCGTACGACTAGATGTGTATAAGAGACAG |
| N501 | AATGATACGGCGACCACCGAGATCTACACTAGATCGCTCGTCGGCAGCGTCAGATGTGTATAAGAGACAG |
| N502 | AATGATACGGCGACCACCGAGATCTACACCTCTCTATTCGTCGGCAGCGTCAGATGTGTATAAGAGACAG |
| N701 | CAAGCAGAAGACGGCATACGAGATTCGCCTTAGTCTCGTGGGCTCGGAGATGTGTATAAGAGACAGCAAGCGTTGGCTTCTCGCATCT |
| N702 | CAAGCAGAAGACGGCATACGAGATCTAGTACGGTCTCGTGGGCTCGGAGATGTGTATAAGAGACAGCAAGCGTTGGCTTCTCGCATCT |
| N703 | CAAGCAGAAGACGGCATACGAGATTTCTGCCTGTCTCGTGGGCTCGGAGATGTGTATAAGAGACAGCAAGCGTTGGCTTCTCGCATCT |
| N704 | CAAGCAGAAGACGGCATACGAGATGCTCAGGAGTCTCGTGGGCTCGGAGATGTGTATAAGAGACAGCAAGCGTTGGCTTCTCGCATCT |
| N705 | CAAGCAGAAGACGGCATACGAGATAGGAGTCCGTCTCGTGGGCTCGGAGATGTGTATAAGAGACAGCAAGCGTTGGCTTCTCGCATCT |
| N706 | CAAGCAGAAGACGGCATACGAGATCATGCCTAGTCTCGTGGGCTCGGAGATGTGTATAAGAGACAGCAAGCGTTGGCTTCTCGCATCT |

**Extended Data Table 2. DNA barcode A sequences.**

| **Sequence Name** | **Sequence (5'-3')** |
| --- | --- |
| Barcode A-1 | /5Phos/AGGCCAGAGCATTCGAACGTGATGTGGCCGATGTTTCG |
| Barcode A-2 | /5Phos/AGGCCAGAGCATTCGAAACATCGGTGGCCGATGTTTCG |
| Barcode A-3 | /5Phos/AGGCCAGAGCATTCGATGCCTAAGTGGCCGATGTTTCG |
| Barcode A-4 | /5Phos/AGGCCAGAGCATTCGAGTGGTCAGTGGCCGATGTTTCG |
| Barcode A-5 | /5Phos/AGGCCAGAGCATTCGACCACTGTGTGGCCGATGTTTCG |
| Barcode A-6 | /5Phos/AGGCCAGAGCATTCGACATTGGCGTGGCCGATGTTTCG |
| Barcode A-7 | /5Phos/AGGCCAGAGCATTCGCAGATCTGGTGGCCGATGTTTCG |
| Barcode A-8 | /5Phos/AGGCCAGAGCATTCGCATCAAGTGTGGCCGATGTTTCG |
| Barcode A-9 | /5Phos/AGGCCAGAGCATTCGCGCTGATCGTGGCCGATGTTTCG |
| Barcode A-10 | /5Phos/AGGCCAGAGCATTCGACAAGCTAGTGGCCGATGTTTCG |
| Barcode A-11 | /5Phos/AGGCCAGAGCATTCGCTGTAGCCGTGGCCGATGTTTCG |
| Barcode A-12 | /5Phos/AGGCCAGAGCATTCGAGTACAAGGTGGCCGATGTTTCG |
| Barcode A-13 | /5Phos/AGGCCAGAGCATTCGAACAACCAGTGGCCGATGTTTCG |
| Barcode A-14 | /5Phos/AGGCCAGAGCATTCGAACCGAGAGTGGCCGATGTTTCG |
| Barcode A-15 | /5Phos/AGGCCAGAGCATTCGAACGCTTAGTGGCCGATGTTTCG |
| Barcode A-16 | /5Phos/AGGCCAGAGCATTCGAAGACGGAGTGGCCGATGTTTCG |
| Barcode A-17 | /5Phos/AGGCCAGAGCATTCGAAGGTACAGTGGCCGATGTTTCG |
| Barcode A-18 | /5Phos/AGGCCAGAGCATTCGACACAGAAGTGGCCGATGTTTCG |
| Barcode A-19 | /5Phos/AGGCCAGAGCATTCGACAGCAGAGTGGCCGATGTTTCG |
| Barcode A-20 | /5Phos/AGGCCAGAGCATTCGACCTCCAAGTGGCCGATGTTTCG |
| Barcode A-21 | /5Phos/AGGCCAGAGCATTCGACGCTCGAGTGGCCGATGTTTCG |
| Barcode A-22 | /5Phos/AGGCCAGAGCATTCGACGTATCAGTGGCCGATGTTTCG |
| Barcode A-23 | /5Phos/AGGCCAGAGCATTCGACTATGCAGTGGCCGATGTTTCG |
| Barcode A-24 | /5Phos/AGGCCAGAGCATTCGAGAGTCAAGTGGCCGATGTTTCG |
| Barcode A-25 | /5Phos/AGGCCAGAGCATTCGAGATCGCAGTGGCCGATGTTTCG |
| Barcode A-26 | /5Phos/AGGCCAGAGCATTCGAGCAGGAAGTGGCCGATGTTTCG |
| Barcode A-27 | /5Phos/AGGCCAGAGCATTCGAGTCACTAGTGGCCGATGTTTCG |
| Barcode A-28 | /5Phos/AGGCCAGAGCATTCGATCCTGTAGTGGCCGATGTTTCG |
| Barcode A-29 | /5Phos/AGGCCAGAGCATTCGATTGAGGAGTGGCCGATGTTTCG |
| Barcode A-30 | /5Phos/AGGCCAGAGCATTCGCAACCACAGTGGCCGATGTTTCG |
| Barcode A-31 | /5Phos/AGGCCAGAGCATTCGGACTAGTAGTGGCCGATGTTTCG |
| Barcode A-32 | /5Phos/AGGCCAGAGCATTCGCAATGGAAGTGGCCGATGTTTCG |
| Barcode A-33 | /5Phos/AGGCCAGAGCATTCGCACTTCGAGTGGCCGATGTTTCG |
| Barcode A-34 | /5Phos/AGGCCAGAGCATTCGCAGCGTTAGTGGCCGATGTTTCG |
| Barcode A-35 | /5Phos/AGGCCAGAGCATTCGCATACCAAGTGGCCGATGTTTCG |
| Barcode A-36 | /5Phos/AGGCCAGAGCATTCGCCAGTTCAGTGGCCGATGTTTCG |
| Barcode A-37 | /5Phos/AGGCCAGAGCATTCGCCGAAGTAGTGGCCGATGTTTCG |
| Barcode A-38 | /5Phos/AGGCCAGAGCATTCGCCGTGAGAGTGGCCGATGTTTCG |
| Barcode A-39 | /5Phos/AGGCCAGAGCATTCGCCTCCTGAGTGGCCGATGTTTCG |
| Barcode A-40 | /5Phos/AGGCCAGAGCATTCGCGAACTTAGTGGCCGATGTTTCG |
| Barcode A-41 | /5Phos/AGGCCAGAGCATTCGCGACTGGAGTGGCCGATGTTTCG |
| Barcode A-42 | /5Phos/AGGCCAGAGCATTCGCGCATACAGTGGCCGATGTTTCG |
| Barcode A-43 | /5Phos/AGGCCAGAGCATTCGCTCAATGAGTGGCCGATGTTTCG |
| Barcode A-44 | /5Phos/AGGCCAGAGCATTCGCTGAGCCAGTGGCCGATGTTTCG |
| Barcode A-45 | /5Phos/AGGCCAGAGCATTCGCTGGCATAGTGGCCGATGTTTCG |
| Barcode A-46 | /5Phos/AGGCCAGAGCATTCGGAATCTGAGTGGCCGATGTTTCG |
| Barcode A-47 | /5Phos/AGGCCAGAGCATTCGCAAGACTAGTGGCCGATGTTTCG |
| Barcode A-48 | /5Phos/AGGCCAGAGCATTCGGAGCTGAAGTGGCCGATGTTTCG |
| Barcode A-49 | /5Phos/AGGCCAGAGCATTCGGATAGACAGTGGCCGATGTTTCG |
| Barcode A-50 | /5Phos/AGGCCAGAGCATTCGGCCACATAGTGGCCGATGTTTCG |

**Extended Data Table 3. DNA barcode B sequences.**

| Sequence Name | Sequence (5'-3') |
| --- | --- |
| barcode B-1 | CAAGCGTTGGCTTCTCGCATCTNNNNNNNNNNAACGTGATATCCACGTGCTTGAG |
| barcode B-2 | CAAGCGTTGGCTTCTCGCATCTNNNNNNNNNNAAACATCGATCCACGTGCTTGAG |
| barcode B-3 | CAAGCGTTGGCTTCTCGCATCTNNNNNNNNNNATGCCTAAATCCACGTGCTTGAG |
| barcode B-4 | CAAGCGTTGGCTTCTCGCATCTNNNNNNNNNNAGTGGTCAATCCACGTGCTTGAG |
| barcode B-5 | CAAGCGTTGGCTTCTCGCATCTNNNNNNNNNNACCACTGTATCCACGTGCTTGAG |
| barcode B-6 | CAAGCGTTGGCTTCTCGCATCTNNNNNNNNNNACATTGGCATCCACGTGCTTGAG |
| barcode B-7 | CAAGCGTTGGCTTCTCGCATCTNNNNNNNNNNCAGATCTGATCCACGTGCTTGAG |
| barcode B-8 | CAAGCGTTGGCTTCTCGCATCTNNNNNNNNNNCATCAAGTATCCACGTGCTTGAG |
| barcode B-9 | CAAGCGTTGGCTTCTCGCATCTNNNNNNNNNNCGCTGATCATCCACGTGCTTGAG |
| barcode B-10 | CAAGCGTTGGCTTCTCGCATCTNNNNNNNNNNACAAGCTAATCCACGTGCTTGAG |
| barcode B-11 | CAAGCGTTGGCTTCTCGCATCTNNNNNNNNNNCTGTAGCCATCCACGTGCTTGAG |
| barcode B-12 | CAAGCGTTGGCTTCTCGCATCTNNNNNNNNNNAGTACAAGATCCACGTGCTTGAG |
| barcode B-13 | CAAGCGTTGGCTTCTCGCATCTNNNNNNNNNNAACAACCAATCCACGTGCTTGAG |
| barcode B-14 | CAAGCGTTGGCTTCTCGCATCTNNNNNNNNNNAACCGAGAATCCACGTGCTTGAG |
| barcode B-15 | CAAGCGTTGGCTTCTCGCATCTNNNNNNNNNNAACGCTTAATCCACGTGCTTGAG |
| barcode B-16 | CAAGCGTTGGCTTCTCGCATCTNNNNNNNNNNAAGACGGAATCCACGTGCTTGAG |
| barcode B-17 | CAAGCGTTGGCTTCTCGCATCTNNNNNNNNNNAAGGTACAATCCACGTGCTTGAG |
| barcode B-18 | CAAGCGTTGGCTTCTCGCATCTNNNNNNNNNNACACAGAAATCCACGTGCTTGAG |
| barcode B-19 | CAAGCGTTGGCTTCTCGCATCTNNNNNNNNNNACAGCAGAATCCACGTGCTTGAG |
| barcode B-20 | CAAGCGTTGGCTTCTCGCATCTNNNNNNNNNNACCTCCAAATCCACGTGCTTGAG |
| barcode B-21 | CAAGCGTTGGCTTCTCGCATCTNNNNNNNNNNACGCTCGAATCCACGTGCTTGAG |
| barcode B-22 | CAAGCGTTGGCTTCTCGCATCTNNNNNNNNNNACGTATCAATCCACGTGCTTGAG |
| barcode B-23 | CAAGCGTTGGCTTCTCGCATCTNNNNNNNNNNACTATGCAATCCACGTGCTTGAG |
| barcode B-24 | CAAGCGTTGGCTTCTCGCATCTNNNNNNNNNNAGAGTCAAATCCACGTGCTTGAG |
| barcode B-25 | CAAGCGTTGGCTTCTCGCATCTNNNNNNNNNNAGATCGCAATCCACGTGCTTGAG |
| barcode B-26 | CAAGCGTTGGCTTCTCGCATCTNNNNNNNNNNAGCAGGAAATCCACGTGCTTGAG |
| barcode B-27 | CAAGCGTTGGCTTCTCGCATCTNNNNNNNNNNAGTCACTAATCCACGTGCTTGAG |
| barcode B-28 | CAAGCGTTGGCTTCTCGCATCTNNNNNNNNNNATCCTGTAATCCACGTGCTTGAG |
| barcode B-29 | CAAGCGTTGGCTTCTCGCATCTNNNNNNNNNNATTGAGGAATCCACGTGCTTGAG |
| barcode B-30 | CAAGCGTTGGCTTCTCGCATCTNNNNNNNNNNCAACCACAATCCACGTGCTTGAG |
| barcode B-31 | CAAGCGTTGGCTTCTCGCATCTNNNNNNNNNNGACTAGTAATCCACGTGCTTGAG |
| barcode B-32 | CAAGCGTTGGCTTCTCGCATCTNNNNNNNNNNCAATGGAAATCCACGTGCTTGAG |
| barcode B-33 | CAAGCGTTGGCTTCTCGCATCTNNNNNNNNNNCACTTCGAATCCACGTGCTTGAG |
| barcode B-34 | CAAGCGTTGGCTTCTCGCATCTNNNNNNNNNNCAGCGTTAATCCACGTGCTTGAG |
| barcode B-35 | CAAGCGTTGGCTTCTCGCATCTNNNNNNNNNNCATACCAAATCCACGTGCTTGAG |
| barcode B-36 | CAAGCGTTGGCTTCTCGCATCTNNNNNNNNNNCCAGTTCAATCCACGTGCTTGAG |
| barcode B-37 | CAAGCGTTGGCTTCTCGCATCTNNNNNNNNNNCCGAAGTAATCCACGTGCTTGAG |
| barcode B-38 | CAAGCGTTGGCTTCTCGCATCTNNNNNNNNNNCCGTGAGAATCCACGTGCTTGAG |
| barcode B-39 | CAAGCGTTGGCTTCTCGCATCTNNNNNNNNNNCCTCCTGAATCCACGTGCTTGAG |
| barcode B-40 | CAAGCGTTGGCTTCTCGCATCTNNNNNNNNNNCGAACTTAATCCACGTGCTTGAG |
| barcode B-41 | CAAGCGTTGGCTTCTCGCATCTNNNNNNNNNNCGACTGGAATCCACGTGCTTGAG |
| barcode B-42 | CAAGCGTTGGCTTCTCGCATCTNNNNNNNNNNCGCATACAATCCACGTGCTTGAG |
| barcode B-43 | CAAGCGTTGGCTTCTCGCATCTNNNNNNNNNNCTCAATGAATCCACGTGCTTGAG |
| barcode B-44 | CAAGCGTTGGCTTCTCGCATCTNNNNNNNNNNCTGAGCCAATCCACGTGCTTGAG |
| barcode B-45 | CAAGCGTTGGCTTCTCGCATCTNNNNNNNNNNCTGGCATAATCCACGTGCTTGAG |
| barcode B-46 | CAAGCGTTGGCTTCTCGCATCTNNNNNNNNNNGAATCTGAATCCACGTGCTTGAG |
| barcode B-47 | CAAGCGTTGGCTTCTCGCATCTNNNNNNNNNNCAAGACTAATCCACGTGCTTGAG |
| barcode B-48 | CAAGCGTTGGCTTCTCGCATCTNNNNNNNNNNGAGCTGAAATCCACGTGCTTGAG |
| barcode B-49 | CAAGCGTTGGCTTCTCGCATCTNNNNNNNNNNGATAGACAATCCACGTGCTTGAG |
| barcode B-50 | CAAGCGTTGGCTTCTCGCATCTNNNNNNNNNNGCCACATAATCCACGTGCTTGAG |

**Extended Data Table 4. Chemicals and reagents used in Spatial ATAC-seq.**

| **Name** | **Catalog number** | **Vender** |
| --- | --- | --- |
| pA-Tn5 Transposase – unloaded | C01070010-20 | Diagenode |
| Tagmentation Buffer (2x) | C01019043-5000 | Diagenode |
| Digitonin | G9441 | Promega |
| 5M NaCl | J60434.AK | Thermo Fisher Scientific |
| Paraformaldehyde solution (16%) | 043368.9M | Thermo Fisher Scientific |
| Glycine | 50046 | Sigma-Aldrich |
| Proteinase K | EO0491 | Thermo Fisher Scientific |
| EDTA (0.5 M) | AM9260G | Thermo Fisher Scientific |
| Fisher Healthcare Tissue-Plus™ O.C.T. Compound | 23-730-571 | Fisher Scientific |
| Triton X-100 | T8787 | Sigma-Aldrich |
| T4 DNA Ligase | M0202L | New England Biolabs |
| T4 DNA Ligase Reaction Buffer | B0202S | New England Biolabs |
| NEBNext High-Fidelity 2X PCR Master Mix | M0541L | New England Biolabs |
| DPBS | 14190144 | Thermo Fisher Scientific |
| DNA Clean & Concentrator-5 | D4014 | Zymo Research |
| dNTP mix | R0192 | Thermo Fisher Scientific |
| EvaGreen Dye 20X in Water | 31000-T | Biotium |
| SPRIselect beads | B23318 | Beckman coulter |
| Poly-L-Lysine Slide | 63478-AS | Election microscopy sciences |
| D5000 ScreenTape | 5067-5588 | Agilent |
| D5000 reagents | 5067-5589 | Agilent |
| Silicon wafer | C04004 | WaferPro |
| Photoresist SU-8 2010 | SU-8 2010 | Microchem Laboratory |
| Tween 20™ 10% | 11332465001 | Sigma-Aldrich |
| Tris-HCl, 1M Solution, pH 8.0 | J22638.K2 | Thermo Fisher Scientific |
| UltraPure™ DNase/RNase-Free Distilled Water | 10977015 | Thermo Fisher Scientific |
| DAPI | 40043 | Biotium |
| Primers, ligation linkers, DNA barcodes | IDT | See Table 1 |
| Chromium Nuclei Isolation kit | 1000493 | 10X Genomics |
